# Deep mutational scanning highlights a new role for cytosolic regions in Hrd1 function

**DOI:** 10.1101/2023.04.03.535444

**Authors:** Brian G. Peterson, Jiwon Hwang, Jennifer E. Russ, Jeremy Schroeder, Peter L. Freddolino, Ryan D. Baldridge

## Abstract

Misfolded endoplasmic reticulum proteins are degraded through a process called endoplasmic reticulum associated degradation (ERAD). Soluble, lumenal ERAD targets are recognized, retrotranslocated across the ER membrane, ubiquitinated, extracted from the membrane, and degraded by the proteasome using an ERAD pathway containing a ubiquitin ligase called Hrd1. To determine how Hrd1 mediates these processes, we developed a deep mutational scanning approach to identify residues involved in Hrd1 function, including those exclusively required for lumenal degradation. We identified several regions required for different Hrd1 functions. Most surprisingly, we found two cytosolic regions of Hrd1 required for lumenal ERAD substrate degradation. Using in vivo and in vitro approaches, we defined roles for disordered regions between structural elements that were required for Hrd1’s ability to autoubiquitinate and interact with substrate. Our results demonstrate that disordered cytosolic regions promote substrate retrotranslocation by controlling Hrd1 activation and establishing directionality of retrotranslocation for lumenal substrate across the endoplasmic reticulum membrane.

## Introduction

Most integral membrane and secretory proteins are translated at the endoplasmic reticulum (ER), where they are folded and undergo protein quality control before distribution to other organelles in the secretory pathway. Newly synthesized proteins that fail to fold are degraded at the proteasome through the conserved pathway called endoplasmic reticulum associated degradation (ERAD). In addition to degradation of misfolded protein, ERAD regulates biosynthetic pathways and degrades key enzymes in sterol synthesis pathways^1–4^. ERAD is critical for maintaining cellular homeostasis and deletion of the ERAD machinery is embryonically lethal in mice^5–9^. When the degradative capacity of the ERAD system is exceeded, unfolded proteins accumulate in the ER and induce the unfolded protein response (UPR) to restore ER proteostasis^10,11^.

Over several decades, genetic and biochemical studies have characterized the ERAD machinery and established fundamental principles for ERAD function. Using *S. cerevisiae* as a model organism, the existence of at least four ERAD pathways have been proposed based on whether the misfolding lesion is in the lumen, membrane, cytosol, or inner nuclear membrane space (ERAD-L, -M, -C, and -INM, respectively)^3,12,13^(for review see^14^). Both ERAD-L and ERAD-M substrates are degraded by the Hrd1 complex normally consisting of Hrd1, Hrd3, Yos9, Der1, and Usa1 (Figure S1A). The Hrd1 complex functions by recruiting substrate to the complex, retrotranslocating lumenal substrate (movement from the ER lumen to the cytosol), followed by substrate ubiquitination, extraction by the Cdc48 complex, and degradation by the proteasome^12,15–18^. Hrd1 is an integral membrane RING-type ubiquitin ligase with eight transmembrane segments that can directly recognize substrates and is required for retrotranslocation^18–22^. Hrd1 forms a heterodimeric retrotranslocation channel with Der1^23,24^. Der1 is a rhomboid pseudo-protease that is essential for ERAD-L under normal physiological conditions^23–25^. Both Hrd1 and Der1 interact directly with Usa1 which scaffolds the heterodimeric-channel interactions and mediates higher-order oligomerization^23,26^. Hrd1 also interacts with Hrd3, a single-pass membrane protein that controls Hrd1 ubiquitination activity, recruits substrates, and bridges an interaction with the substrate-recruiting protein called Yos9^12,27–29^. The Hrd1 complex is proposed to be dynamic, with structural studies finding Hrd1 as a “monomeric” (Der1-Hrd1-Hrd3-Usa1) or dimeric (Hrd3-Hrd1-Hrd1-Hrd3) complex state^19,23,30,31^.

Genetic, biochemical, and structural studies support a central role for Hrd1 in ERAD. Overexpression of Hrd1 bypasses the requirement for other complex components in vivo^20^ and Hrd1 can independently recognize, retrotranslocate, and ubiquitinate substrate in vitro^18,21,22^. Moreover, Hrd1 controls the activity of the complex by activating itself through autoubiquitination within its RING domain^21^. It is currently unclear how Hrd1 autoubiquitination results in Hrd1 activity; proposals include that autoubiquitination drives a conformational change in Hrd1 that permits substrate retrotranslocation^21,22^, exposes a cytosolic substrate binding site^22^, or alters the oligomeric state from a regulatory to an active stoichiometry^30^.

To dissect the mechanics of Hrd1 function, we developed fluorescent reporter substrates and used unbiased, deep mutational scanning (DMS) to identify residues within Hrd1 required for ERAD. Our approach highlighted two surprising regions on the cytosolic face of Hrd1 that were specifically required for degradation of lumenal ERAD substrates (ERAD-L). These important cytosolic regions are predicted to be disordered but have intrinsic elements that were required for their function in ERAD. Using both in vivo and in vitro assays, we found that the first disordered cytosolic loop between transmembrane segments 6 and 7 is required for Hrd1 autoubiquitination. The second disordered region falls within the C-terminal region and forms a cytosolic substrate binding domain required to promote retrotranslocation of lumenal substrates across the membrane. Our work unveils new mechanics for lumenal substrate selection and retrotranslocation across the ER membrane.

## Results

### Deep mutational scanning of the Hrd1 ubiquitin ligase

We developed an unbiased, deep mutational scanning approach to dissect Hrd1 function. We designed fluorophore-based reporters using model ERAD substrates to allow for high-throughput screening by flow cytometry (Figures 1A, and S1B-S1D)^1,32–35^. For our assay, we selected CPY* (GFP-CPY*) and Hmg2 (Hmg2-RFP), as model ERAD-L and ERAD-M substrates^1,33^. We observed clear fluorescence shifts between wild-type Hrd1(WT) and the non-functional RING domain mutant Hrd1(C399S) (Figure 1A and S1D). Using flow cytometry, we followed substrate degradation by allowing cells to enter stationary phase, which reduces new translation of ERAD substrates (hereafter called a saturated chase) (Figures 1B and S1E)^15^. We reasoned this method would give us better recovery of sorted cells compared to the more widely-used cycloheximide chase^1,36^.

**Figure 1.**
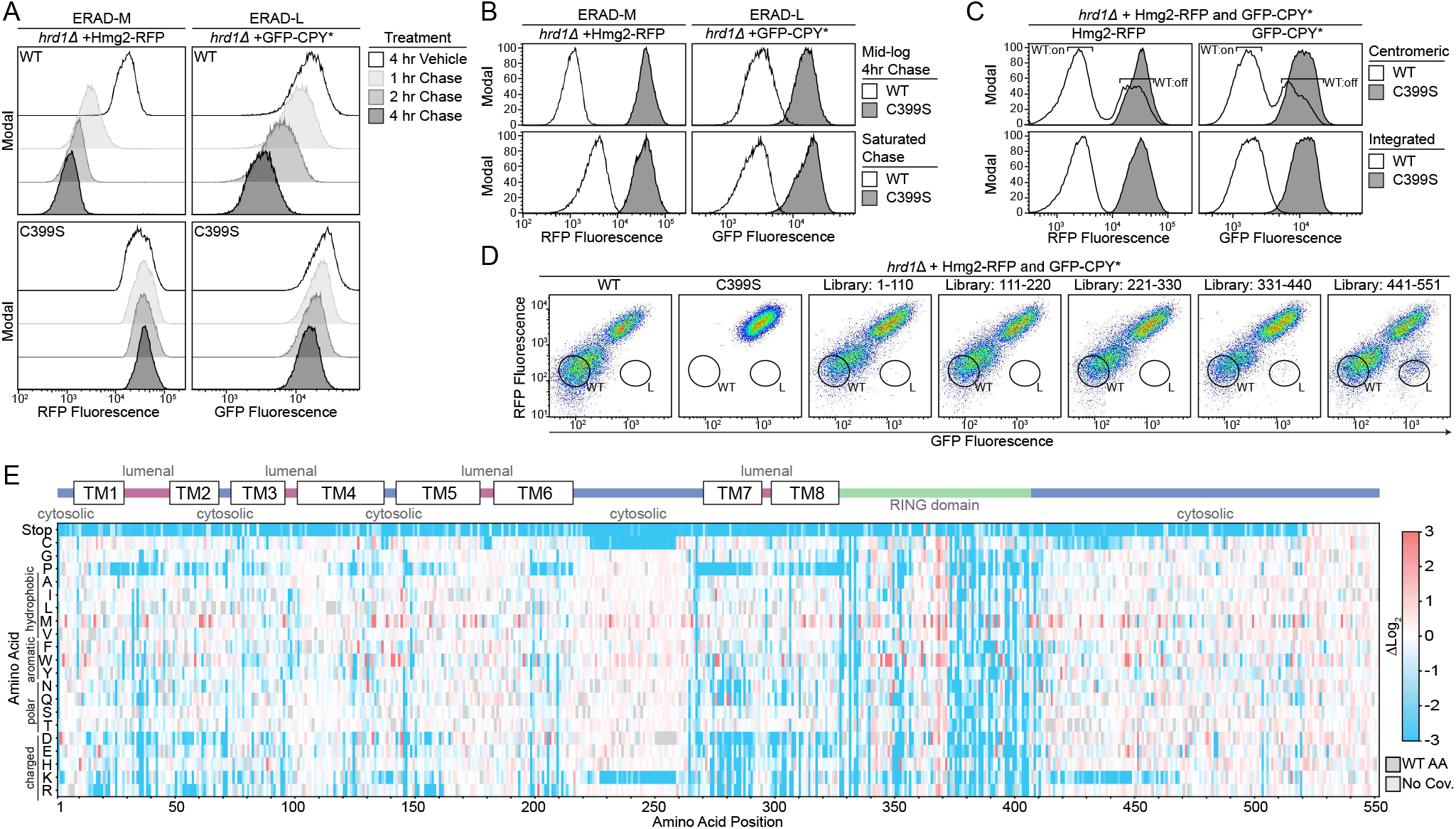
Deep mutational scanning of the Hrd1 ubiquitin ligase. A) Degradation of Hmg2-RFP and GFP-CPY* were followed using flow cytometry after the addition of either 0.1% ethanol (vehicle control), 10 µg/mL zaragozic acid (for Hmg2 chase), or 50 µg/mL cycloheximide (for CPY* chase). Experiments were performed in *hrd1Δ* cells expressing either wild-type Hrd1(WT) or a non-functional Hrd1(C399S). Histograms were scaled as a percentage of maximum cell count (Modal). B) Substrate degradation was followed using flow cytometry during either mid-log phase growth treated as in (A) (Mid-log Chase, top panels) or with cells grown to saturation and no pharmacological treatment (Saturated Chase, bottom panels) C) As in (A) except with *hrd1Δ* cells co-expressing Hmg2-RFP and GFP-CPY*. Cells were complemented by either centromeric Hrd1 plasmids or with Hrd1 integration and subjected to a saturated chase. Centromeric Hrd1(WT) forms two populations of expression: either expressing (“WT:on”) or low expression (“WT:off”) as observed previously^39^. D) Fluorescence-activated cell sorting (FACS) analysis of *hrd1Δ* cells expressing Hmg2-RFP (y-axis) and GFP-CPY* (x-axis) were transformed with centromeric plasmids containing either wild-type Hrd1(WT), inactive Hrd1(C399S), or one of five different Hrd1 mutant libraries. Wildtype-like (WT) and ERAD-L defective (L) cells were sorted into bins as indicated and collected for downstream analysis. E) Top: Topology diagram of Hrd1 with transmembrane segments displayed as TM1-8. Colors indicate the cytosolic (blue), lumenal (magenta), and cytosolic RING domain (green). Bottom: Deep mutational scanning results of cells sorted from the wildtype-like bins in (D) were displayed as a heatmap showing single codon changes that were enriched (red) or depleted (blue) compared to the input library. Individual amino acids are on the y-axis, and the Hrd1 amino acid position is on the x-axis. Dark gray boxes indicate the wildtype amino acid and light gray boxes indicate a lack of sequencing coverage.

We used tiling primer mutagenesis to mutagenize five subregions (approximately 110 amino acids each) that span Hrd1 and are compatible with short-read Illumina sequencing^37^. We inserted the mutagenized DNA fragments into a centromeric plasmid backbone using in-cell homologous recombination^38^. While expression from centromeric plasmids switches between on- and off-expression states^39^, we opted for centromeric plasmids, rather than genomic integration, to increase transformation efficiency (Figure 1C and Table S1). We subjected these cells to a saturated chase and isolated wildtype-like (WT) and ERAD-L defective (L) populations using fluorescence-activated cell sorting (FACS) (Figure 1D). After outgrowth, we confirmed the phenotype of the sorted populations, followed by DNA extraction, library preparation, Illumina sequencing, and analysis (Figure S1F and S1G).

We began our analysis with wildtype-like sorted cells (WT bin, Figure 1D) with only single amino acid changes (covering 99.7% of the possible substitutions). We compared the relative ratio of mutations in the WT bin to our input libraries and visualized the results with a heatmap (Figure 1E). To validate our screening and analysis pipeline, we considered mutations expected to prevent wild-type function based on prior knowledge. First, we observed stop codons were strongly depleted (blue in heatmap) in our WT bin, except within the last 30 amino acids of Hrd1 (Figure 1E; Table S2-S5). Second, we expected that prolines would disrupt the transmembrane segments and prevent normal function^40,41^. Indeed, we observed a strong depletion of prolines in most transmembrane segments. Finally, we expected that mutations in the RING domain of Hrd1 would prevent function. As expected, mutations in the RING-finger cross-brace motifs were highly depleted for most substitutions. Together, these results demonstrated that our screening and analysis pipelines performed as expected.

### Transmembrane segments 1 and 2 control complex specificity through distinct mechanics

To specifically identify portions of Hrd1 contributing to ERAD-L, we shifted our attention to cells that were sorted as defective in CPY* degradation (ERAD-L), but functional in Hmg2 degradation (ERAD-M) (L bin, Figures 1D and S1G). We looked for enriched single-point mutations and focused on results with low false discovery rates (FDR) (Figures 2A and S2A-S2C, Table S5-S7). We observed clear enrichment of mutations that prevented ERAD-L substrate degradation scattered across transmembrane segments 1 and 2 and in the C-terminal region following the RING domain.

**Figure 2.**
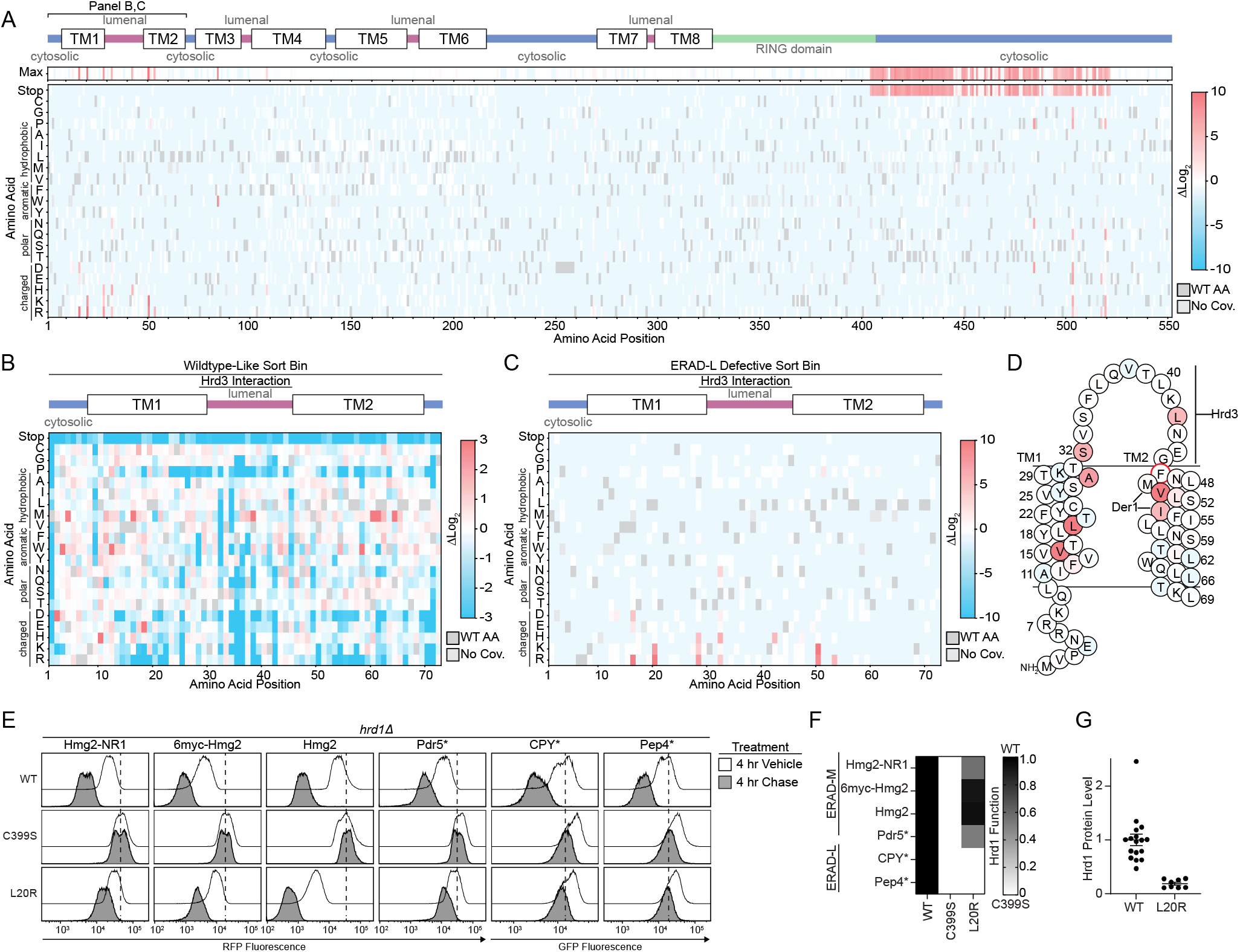
Transmembrane segments 1 and 2 control complex specificity through distinct mechanics. A) Top: Topology diagram of Hrd1 with transmembrane segments shown as TM1-8. Colors indicate the cytosolic (blue), lumenal (magenta), and cytosolic RING domain (green). Middle: Heatmap showing the highest enrichment value at each amino acid position from cells sorted into the ERAD-L defective bin. Bottom: Deep mutational scanning results of cells sorted into the ERAD-L defective bins displayed as a heatmap showing single codon changes that were enriched (red) or depleted (blue) compared to the input library. Color transparency was adjusted based on false discovery rates (FDR). FDR below 0.1% were set to 0% transparent and FDR values between 0.1% to 100% were used to adjust transparency from 0% (opaque) to 90% transparent (see also Figures S2A-S2C). Individual amino acids are on the y-axis, and the Hrd1 amino acid position is on the x-axis. Dark gray boxes indicate the wildtype amino acid and light gray boxes indicate lack of coverage. B) As in (A), but with a heatmap representing the cells sorted into the wildtype-like bin between residues Met1-Gly72. Transparency was not adjusted based on FDR. C) As in (A) but showing only amino acids Met1-Gly72. D) Topology diagram highlighting positions of ERAD-L defective Hrd1 mutations from (C), with the color scheme matching enrichment values. The red circle (F46R) indicates an ERAD-L defective mutant isolated during screen development. E) Flow cytometry analysis of *hrd1Δ* cells expressing the indicated ERAD substrates and Hrd1(WT), inactive Hrd1(C399S), or Hrd1 variants. Cells were treated for 4 hours with 0.1% ethanol (vehicle, solid black line with no fill) or chased for 4 hours with cycloheximide or zaragozic acid (solid black line with gray fill). The dashed line highlights the position of Hrd1(C399S) after a 4-hour chase. F) Quantification of (E). Hrd1(WT) is set to 1 (full function, black) and inactive Hrd1(C399S) is set to 0 (no function, white). Values outside of the range were set to 0 or 1 (see also Figure S2D). G) Protein levels of Hrd1 or Hrd1(L20R) expressed from native promoters on centromeric plasmids were determined by quantification of immunoblots normalized to wild-type Hrd1(WT) levels displayed as mean +/− SEM. For this figure, the number of quantified replicates and individual values are shown in Table S8 and S9.

The Hrd1 lumenal segment between transmembrane segments 1 and 2 was previously reported to interact with Hrd3^19,23,27^. Our screening data highlights the importance of this interaction interface because mutations to many amino acids including polar, charged, or proline were not permissive to Hrd1 function, presumably through loss of Hrd3 association (Figure 2B). Notably, we found specific mutations clustered near or within the Hrd3 interaction site (Ala28, Ser32, Leu42, Phe46) that reduced ERAD-L function (Figures 2C, 2D, and S2D).

We also identified mutations clustering within transmembrane segments 1 (Val16, Leu20) and 2 (Val50, Ile53) that specifically prevented degradation of ERAD-L substrates (Figures 2C, 2D). Based on earlier studies, Met49 and Ile53 are likely to be positioned closely to Der1^23^, a component required for ERAD-L degradation^25^. We expect that mutations in Val50 and Ile53 are likely to perturb Hrd1/Der1 interaction, explaining their ERAD-L defects. However, residues within transmembrane segment 1 have not been shown to interact with Der1, so it is unlikely Val16 and Leu20 substitutions disrupt Hrd1/Der1 interactions^23^. To study this region, we selected the highest-enriched variant from transmembrane segment 1, Hrd1(L20R) (Figures 2C and 2D). We confirmed Hrd1(L20R) was unable to degrade either ERAD-L substrate (GFP-CPY* and GFP-Pep4*) but still degraded all ERAD-M substrates (Hmg2-NR1, 6myc-Hmg2, Hmg2, and Pdr5*), thus demonstrating a strong and specific defect for ERAD-L (Figure 2E, 2F, and S2D). Curiously, Hrd1(L20R) had enhanced degradation activity for Hmg2, which was apparent because of reduced steady-state levels in vehicle-treated cells (Figure 2E). We found Hrd1(L20R) expression was around 1/5th of Hrd1(WT), eliminating the possibility that elevated levels of Hrd1(L20R) enhanced Hmg2 degradation (Figure 2G). In summary, we found clusters of mutations across transmembrane segments 1 and 2 that likely disrupt ERAD-L in different ways: preventing Hrd3 interaction, Der1 interaction, and altering Hrd1 specificity.

### Usa1 interacts with the Hrd1 C-terminal region

Our screening results also demonstrated that the Hrd1 C-terminal region was required for degradation of ERAD-L, but not ERAD-M, substrates (Figure 2A). We observed an enrichment of stop codons beginning at residue Phe404 and ending at Ile521 that prevented ERAD-L degradation (Figures 3A, S2A-S2C). We confirmed that the complete removal of the C-terminal region (Hrd1(Δ408-551)) prevented ERAD-L substrate degradation but maintained the ability to degrade all four ERAD-M substrates (Figures 3B and S3A).

**Figure 3.**
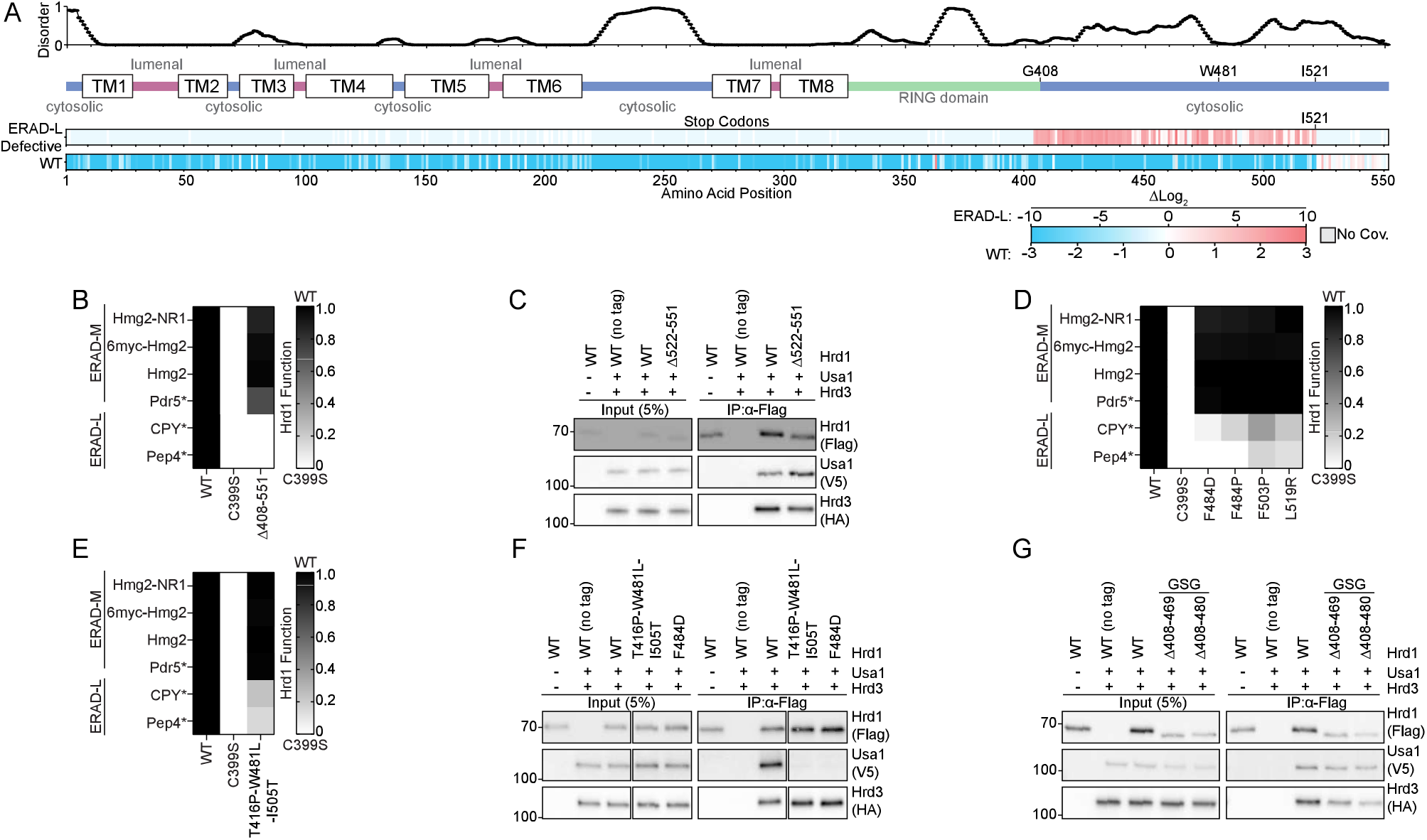
Usa1 interacts with the Hrd1 C-terminal region. A) Top: Line chart representing predicted disorder (PONDR VLXT^57^) of Hrd1 normalized 0 (ordered) to 1 (disordered). Middle: Topology diagram of Hrd1 with transmembrane segments shown as TM1-8. Bottom: deep mutational scanning results of stop codon enrichment values within the ERAD-L defective sorting bin (ERAD-L Defective) and wildtype-like sort bin (WT). Amino acid position is on the x-axis. Missing sequencing coverage is shown in light gray. The ERAD-L defective heatmap’s color transparency was adjusted based on FDR. FDR below 0.1% were set to 0% transparent and FDR values between 0.1% to 100% were used to adjust transparency from 0% (opaque) to 90% transparent. Transparency was not adjusted using FDR for the wildtype-like heatmap. B) Degradation of ERAD substrates were followed by flow cytometry and summarized in a heatmap. The indicated Hrd1 variants were integrated in a *hrd1Δ* expressing individual ERAD substrates and subjected to a 4-hour mid-log chase. Wild-type Hrd1(WT) is set to 1 (full function, black) and inactive Hrd1(C399S) is set to 0 (no function, white). Values outside of the range were set to 0 or 1 (see also Figure S3A). C) Co-immunoprecipitation of the Hrd1 complex was performed with the indicated Hrd1 variants. 3xHA-Hrd3, 3xV5-Usa1, and Hrd1-3xFlag were integrated into *hrd1Δhrd3Δusa1Δ* cells, lysed, and immunoprecipitated with anti-Flag antibodies. Input represents 5% of the cleared lysate. This immunoblot is representative of 3 independent replicates. D) As in (B) with the indicated Hrd1 variants. (See Figure S3B). E) As in (B) with the indicated Hrd1 variants. F) As in (C) with the indicated Hrd1 variants. G) As in (C) with the indicated Hrd1 variants. GSG refers to a poly-Gly-Ser-Gly linker the same length as indicated deletion. For this figure, the number of quantified replicates and individual values are shown in Table S8.

One possible explanation for this result is that loss of the C-terminal region of Hrd1 prevents an interaction with Usa1. Usa1 is required for ERAD-L substrate degradation and the C-terminal region of Hrd1 (Lys518-Ile551) was previously demonstrated to interact with Usa1^12,26^. Correspondingly, our screen identified a point mutation within the reported Usa1 interaction site at Leu519 that was defective in degradation of ERAD-L substrates (Figure 2A and Table S5-S7). However, in our experiments, truncations beginning at Glu522 retained normal ERAD-L function, meaning that residues after Ile521 were not required for interaction with Usa1 (Figure 3A). Using co-immunoprecipitation we confirmed that Hrd1(Δ522-551) maintained normal Hrd1/Usa1 interaction, refining the Hrd1/Usa1 interaction site to end at Ile521 (Figure 3C).

Based on the deep mutational scanning data, we identified non-truncation point mutations between Phe484 to Leu519 that were enriched in the ERAD-L defective populations (Figure 2A and Table S5-S7). We confirmed that these point mutations were defective in ERAD-L, but not ERAD-M, substrate degradation (Figure 3D, S3B). In addition, during screening development, we discovered an ERAD-L defective triple point mutant within this region, Hrd1(T416P,W481L,I505T), and confirmed the ERAD-L defective phenotype (Figure 3E). For each of these mutants, the steady-state levels were similar to wild-type Hrd1, indicating the ERAD-L defects were not related to the protein stability (Figure S3C). Based on the proximity of these residues to the reported Hrd1/Usa1 interface, we suspected these point mutations would disrupt Hrd1/Usa1 interaction^26^. Co-immunoprecipitation experiments confirmed that Hrd1(T416P,W481L,I505T) and Hrd1(F484D) failed to interact with Usa1, but interacted normally with Hrd3 (Figure 3F).

Given the observed enrichment of stop codons starting at Phe404, we wondered whether the region between the RING-domain and Trp481 was also important for Usa1 interaction. To determine whether Gly408-Thr480 was important for Usa1 interaction, we replaced these residues with a poly-Gly-Ser-Gly linker of the same length (Hrd1(Δ408-480_GSG)) and a slightly shorter replacement region from Gly408-Met469 (Hrd1(Δ408-469_GSG)). Using co-immunoprecipitation, we found that both Hrd1(Δ408-480_GSG) and Hrd1(Δ408-469_GSG) maintained normal Usa1 and Hrd3 interactions (Figure 3G). These data allowed us to refine the Hrd1/Usa1 interaction interface to Hrd1 Trp481-Ile521, which is required for ERAD-L, but not ERAD-M.

### A disordered C-terminal region is required for retrotranslocation

The requirement of a cytosolic region (Gly408-Thr480) for lumenal substrate degradation was surprising because Hrd1(Δ408-480_GSG) appeared to interact with the other ERAD components properly. In addition, this region was predicted to be disordered (Figure 3A) so we considered whether it would function solely as a spacer; in this case, enrichment of stop codons could be a function of removing the subsequent Usa1 interaction interface. We reasoned that if Gly408-Thr480 served strictly as a spacer, any replacement sequence would maintain wildtype-like activity. First, we tested Hrd1(Δ408-469_GSG) and Hrd1(Δ408-480_GSG), finding both constructs were completely unable to degrade ERAD-L substrates but were still able to degrade ERAD-M substrates (Figure 4A). Next, we replaced Hrd1 Gly408-Met469 with the corresponding regions of Hrd1 from *Saccharomyces kudriavzevii* (Hrd1(*Sk*)) and *Lachancea nothofagi* (Hrd1(*Ln*)) (Figure S4A). Each chimera was able to degrade ERAD-L and ERAD-M substrates, although less efficiently than endogenous *S. cerevisiae* Hrd1(WT) (Figure 4A). We measured steady-state Hrd1 levels and found each construct exhibited reduced Hrd1 stability (Figures 4B and 4C). However, stability alone did not explain the activity defects because Hrd1(*Sk*), Hrd1(Δ408-469_GSG), and Hrd1(Δ408-480_GSG) had similar stability but disparate degradation capability (Figures 4A-4C). These data support the idea that the amino acid composition of the disordered Gly408-Thr480 region is important for Hrd1 function in degradation of ERAD-L substrates.

**Figure 4.**
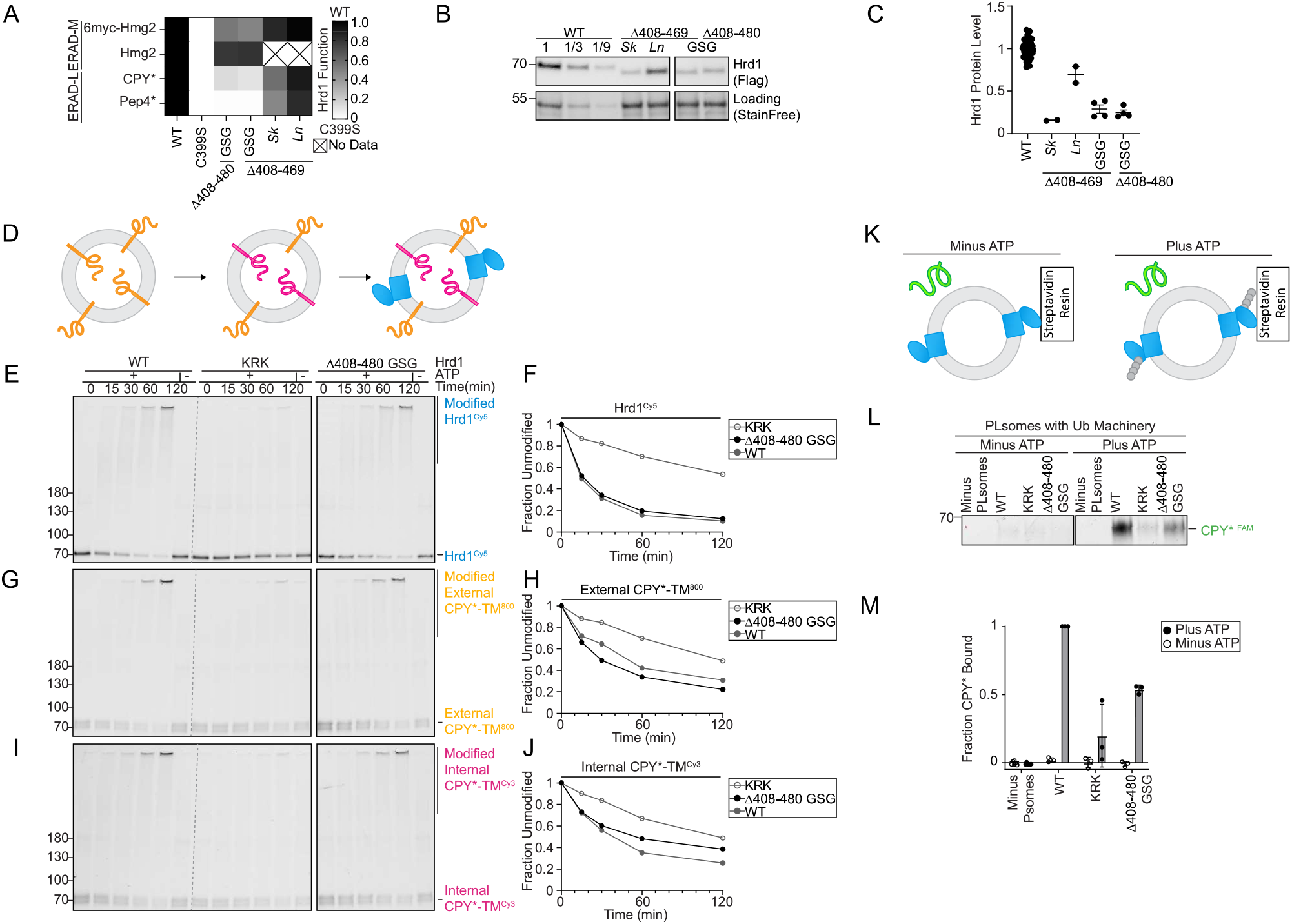
A disordered C-terminal region is required for retrotranslocation. A) Degradation of ERAD substrates by individual Hrd1 variants were followed by flow cytometry and summarized in a heatmap. The indicated Hrd1 variants were integrated in a *hrd1Δ* expressing individual ERAD substrates and subjected to a 4-hour mid-log chase. For “GSG” Hrd1 samples, the region indicated was replaced with a poly-Gly-Ser-Gly linker of the same length as indicated deletion. *Sk* (*S. kudriavzevii*) and *Ln* (*L. nothofagi*) are chimeras of *S. cerevisiae* Hrd1 replaced with homologous regions from indicated species (see Figure S4A). Wild-type Hrd1(WT) is set to 1 (full function, black) and inactive Hrd1(C399S) is set to 0 (no function, white). B) Expression levels of Hrd1 variants were determined by immunoblotting. *hrd1Δ* cells expressing Hrd1-3xFlag constructs from (A). The first three lanes are a calibration curve generated with a dilution of the wild-type Hrd1 lysate. Total protein was visualized by stain-free technology as a loading control. C) Quantification of (B), normalized to wild-type Hrd1 expression and displayed as the mean +/− SEM. D) Schematic of the in vitro retrotranslocation assay. CPY*-TM labeled C-terminally with DyLight^800^ (orange) was reconstituted into proteoliposomes. The C-terminal DyLight^800^ peptide of internally-oriented substrate was exchanged for a new peptide containing Cy3 (magenta) using sortase A followed by incorporation of Hrd1^Cy5^ (blue). E) In vitro autoubiquitination of Hrd1^Cy5^ (blue) in a reconstituted proteoliposome system with externally-oriented CPY*-TM^800^ (orange), internally-oriented CPY*-TM^Cy3^ (magenta). Wild-type Hrd1 (WT-positive control), a retrotranslocation-defective Hrd1 (Hrd1(KRK)-negative control), or Hrd1(*Δ*408-480 GSG) were reconstituted and incubated with recombinant ubiquitination machinery for the indicated times. Samples were analyzed by SDS-PAGE and in-gel fluorescence scanning to visualize Hrd1. Red pixels indicate saturation of signal during the imaging. For second and third replicate see (Figures S4B-S4M). F) Quantification of (E), the fraction of unmodified Hrd1^Cy^^5^. G) As in (E) showing external CPY*-TM^800^. H) Quantification of (G), the unmodified external CPY*-TM^800^. I) As in (E) showing internal CPY*-TM^Cy^^3^. J) Quantification of (I), the unmodified internal CPY*-TM^Cy^^3^. K) Schematic for soluble CPY* (green) interaction assay in proteoliposomes. Hrd1 is shown in blue and ubiquitin chains in gray. L) Hrd1 proteoliposomes (PLsomes) were immobilized at 1 µM and incubated with recombinant ubiquitination machinery, +/− ATP, washed, and incubated with 100nM CPY*^FAM^ (green) (see K). Samples were eluted with biotin and analyzed by SDS-PAGE and in-gel fluorescence scanning. Red pixels indicate image saturation. M) Quantification of (L) shown as the mean +/− SEM, from three independent experiments. For this figure, the number of quantified replicates and individual values are shown in Table S8 and S9.

To identify the specific function of this disordered region, we turned to an in vitro reconstituted system enabling dissection of the individual steps in the ERAD process. Hrd1 was reconstituted into proteoliposomes with an ERAD-L substrate (CPY*-TM) that was C-terminally labeled with an organic fluorophore to track CPY*-TM orientation (Figure 4D)^21^. Hrd1 function was followed in different ways. First, by Hrd1 autoubiquitination (Figures 4E and 4F); second, by ubiquitination of externally-oriented substrate (active ubiquitin ligase activity) (Figures 4G and 4H); third, by ubiquitination of internally-orientated substrate (demonstration of substrate retrotranslocation) (Figures 4I and 4J). As controls for this assay, we used wild-type Hrd1 and a previously-established retrotranslocation-defective Hrd1 (Hrd1(KRK))^21,22^. Hrd1 and Hrd1(KRK) performed as expected (Figures 4E-4J). Hrd1(Δ408-480_GSG) autoubiquitination was similar to wild-type Hrd1 (Figure 4E and 4F) as was the ubiquitination of externally-oriented substrate (Figures 4G and 4H). However, this Hrd1 variant showed a notably reduced ability to retrotranslocate and ubiquitinate internally-oriented substrate (Figures 4I and 4J). This indicates Hrd1(Δ408-480_GSG) retrotranslocation function is partially impaired, indicating that the cytosolic-facing Gly408-Thr480 disordered region of Hrd1 plays an important role in substrate retrotranslocation, specifically.

Hrd1 was previously proposed to contain a cytosolic substrate binding site that becomes exposed following autoubiquitination^22^. To determine whether Gly408-Thr480 directly interacts with substrates, we used a previously described proteoliposome-based substrate interaction assay^22^. We reconstituted Hrd1 into proteoliposomes and immobilized the proteoliposomes with an affinity resin. We incubated the proteoliposomes with purified ubiquitination machinery either in the presence, or absence of ATP. After removing the ubiquitination machinery, we incubated the proteoliposomes with a soluble ERAD-L substrate (CPY*) and followed CPY* interaction with the proteoliposomes (Figure 4K). As expected, Hrd1(WT) efficiently interacted with CPY* after autoubiquitination, while Hrd1(KRK) did not (Figures 4L and 4M)^22^. Hrd1(Δ408-480_GSG) was unable to efficiently interact with CPY* following autoubiquitination, even though the autoubiquitination activity was comparable to wild-type Hrd1 (Figure 4E). Taken together, these results support a role for a previously-overlooked disordered C-terminal region in Hrd1 (Gly408-Thr480). This region is necessary for efficient ERAD-L substrate retrotranslocation through direct substrate interaction.

### Hrd1 autoubiquitination outside of the RING domain restricts function

We noticed that the C-terminal disordered region was largely devoid of lysine and cysteine residues (Figure S4A). This observation was complimented by our screening data that showed mutations adding lysine or cysteine residues were largely depleted between Gly408-Met469 in the wildtype-like sorted group (Figures 1E and 5B). We directly tested this observation by substituting lysine across the C-terminal region (Gly408-Ser475) and assaying ERAD function. Consistent with our screening data, lysine substitutions within Gly408-Met469, largely reduced activity against both classes of substrates, but demonstrated no change in specificity between ERAD-L and ERAD-M substrates (Figures 5C and S5A).

**Figure 5.**
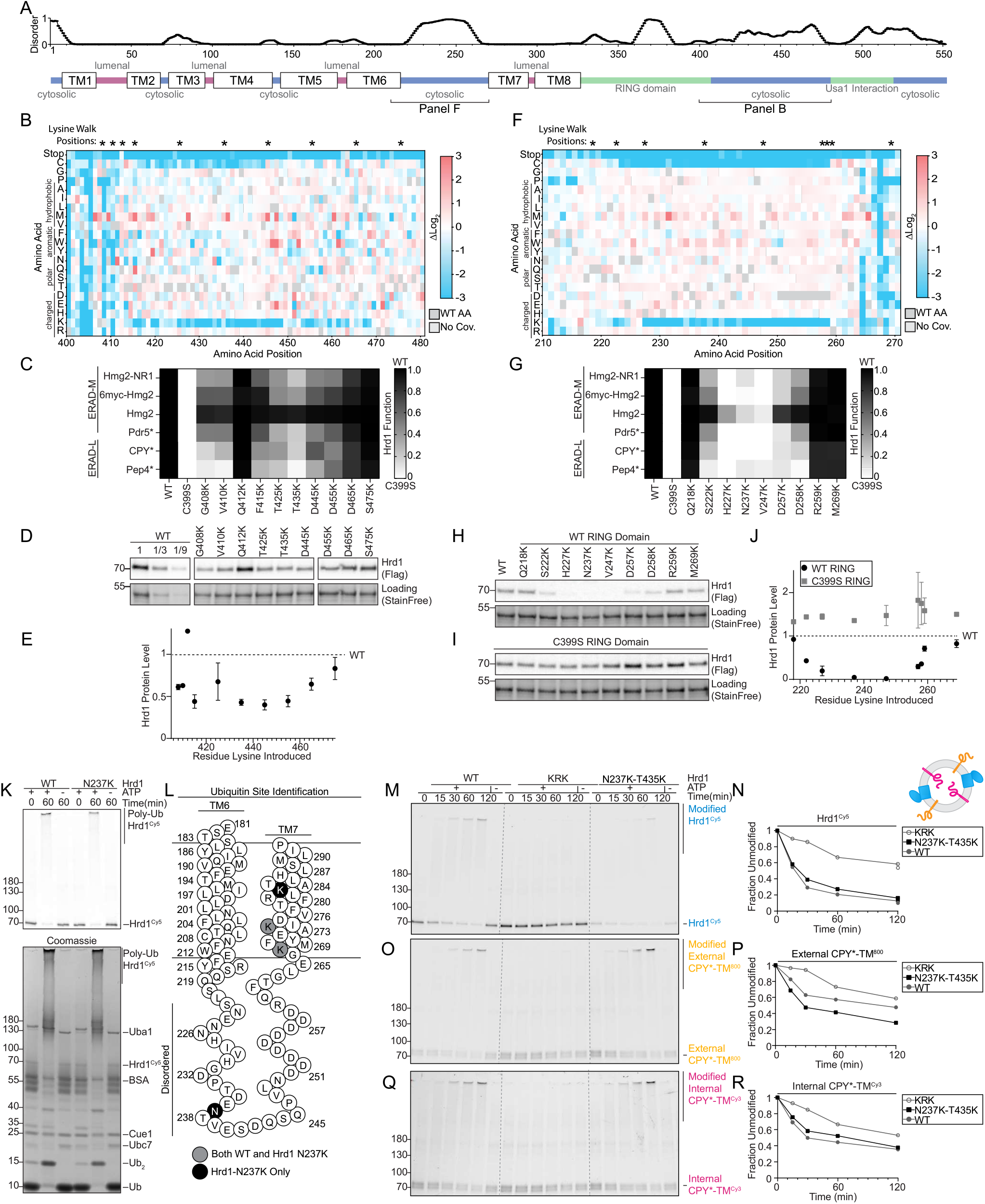
Disordered cytosolic regions are devoid of lysine and cysteine. A) Top: Line chart representing predicted disorder (PONDR VLXT^57^) of Hrd1 normalized 0 (ordered) to 1 (disordered). Bottom: Topology diagram of Hrd1 with transmembrane segments shown as TM1-8. B) Top: “*” indicate positions where lysine is substituted in panels C, D, and E. Bottom: heatmap representing the deep mutational scanning results of Hrd1 variants sorted into the wildtype-like bin. Amino acids from Arg400 to Thr480 are shown. Enrichment values are compared to the input library. Dark gray boxes indicate the wildtype amino acid and light gray boxes indicate lack of coverage. C) Degradation of ERAD substrates by individual Hrd1 variants were followed by flow cytometry and summarized in a heatmap. The indicated Hrd1 variants were integrated in a *hrd1Δ* expressing individual ERAD substrates and subjected to a 4-hour mid-log chase. Wild-type Hrd1(WT) is set to 1 (full function, black) and inactive Hrd1(C399S) is set to 0 (no function, white). Values outside of this range were set to 0 or 1 (see Figure S5A). D) Expression levels of Hrd1 variants were determined by immunoblotting. *hrd1Δ* cells expressing Hrd1-3xFlag constructs from (C). Total protein was visualized by stain-free technology as a loading control. The first three lanes are a calibration curve generated with a dilution of the wild-type Hrd1 lysate. E) Quantification of (D), normalized to wild-type Hrd1 expression and displayed as the mean +/− SEM. (See Figure S5B). F) As in (B), but for amino acid positions Asn210-Tyr270. G) As in (C), but with the indicated Hrd1 variants. (See Figure S5C). H) As in (D), but with the indicated Hrd1 variants. I) As in (H), but containing E3-inactivating C399S RING mutation. J) Quantification of (H) as black filled circles and (I) gray filled squares, normalized to wild-type Hrd1 (black dashed line). (See Figure S5E and S5F). K) Purified Hrd1(WT) or Hrd1(N237K) was incubated with recombinant ubiquitination machinery. Samples were separated by SDS-PAGE, and poly-ubiquitinated Hrd1 bands were excised from the gel and sent for mass spectrometry-based identification of ubiquitination sites. Top: Hrd1^Cy5^ fluorescence scanning of the in vitro ubiquitination reactions. Bottom: Coomassie staining. L) Topology diagram showing TM6 and TM7 and the predicted disordered intervening cytosolic loop. Lysines that were identified as ubiquitinated for both wild-type Hrd1 and Hrd1(N237K) are displayed as gray filled circles. Lysines ubiquitinated only in Hrd1(N237K) are displayed as a black filled circles. M) In vitro autoubiquitination of Hrd1^Cy5^ (blue) in a reconstituted proteoliposome system with externally-oriented CPY*-TM^800^ (orange), internally-oriented CPY*-TM^Cy3^ (magenta). Wild-type Hrd1 (WT-positive control), a retrotranslocation-defective Hrd1 (Hrd1(KRK)-negative control), or Hrd1(N237K-T435K) were reconstituted and incubated with recombinant ubiquitination machinery for the indicated times. Samples were analyzed by SDS-PAGE and in-gel fluorescence scanning to visualize Hrd1. Red pixels indicate saturation of signal during the imaging. For second and third replicate see (Figures S5I-S5T). N) Quantification of (M), showing unmodified Hrd1^Cy^^5^. O) As in (M) showing external CPY*-TM^800^. P) Quantification of (O), showing unmodified external CPY*-TM^800^. Q) As in (M) showing internal CPY*-TM^Cy^^3^. R) Quantification (Q), showing unmodified internal CPY*-TM^Cy^^3^. For this figure, the number of quantified replicates and individual values are shown in Table S8 and S9.

We suspected that the depletion of lysine residues in proximity to the RING domain could stem from one of three possibilities. First, lysines could reduce Hrd1 function because they are ubiquitinated, resulting in Hrd1 degradation. Alternatively, lysine ubiquitination could sterically obstruct the observed substrate interaction with this C-terminal region. Lastly, ubiquitination in this C-terminal region could sterically inhibit substrate passage through the retrotranslocon. We determined whether lysine substitutions within Hrd1 residues Gly408-Met469 were destabilizing by measuring Hrd1 steady-state levels. We found that lysine substitutions at all tested residues, except Gln412, resulted in reduced Hrd1 levels (Figures 5D, 5E, and S5B), roughly corresponding to each mutant’s overall degradation function (Figure 5C-5E, S5A-S5B, and S5H). These data support the hypothesis that lysine and cysteine residues were not tolerated in the Gly408-Met469 region because they were ubiquitinated resulting in Hrd1 turnover rather than interfering with substrate interaction or retrotranslocation directly.

We observed a second region on the cytosolic face that had similar characteristics to the C-terminal region we implicated in direct substrate interaction (Gly408-Thr480). Both regions were predicted to be disordered, devoid of lysine, and devoid of cysteine. This second region falls within a cytosolic loop between transmembrane segments 6 and 7 (Ser222-Asp258, Figure 1E, 5A, 5F). Similar to the Gly408-Met469 region, our functional screen found that Ser222-Asp258 was tolerant to most substitutions, except for lysine or cysteine (Figure 5F). As with the C-terminal results, substituting lysine or cysteine within Ser222-Asp258 resulted in severe defects degrading ERAD-M and ERAD-L substrates (Figures 5G, S5C, and S5D). Likewise, we found that steady-state levels of Hrd1 variants with lysine or cysteine mutations within Ser222-Asp258 were reduced (Figures 5H, 5J, S5E, and S5F) and this destabilization correlated with the severity of Hrd1 function loss (Figure 5G, 5H, 5J, S5C-S5F, S5H). Destabilization of Hrd1 with substitutions within Ser222-Asp258 was caused by Hrd1 autoubiquitination, rather than another ubiquitin ligase, as the same lysine substitutions in an inactive Hrd1(C399S) mutant did not lead to destabilization (Figures 5I,5J and S5G). Together, our data suggest that lysine or cysteine substitutions between positions Ser222-Asp258 destabilized Hrd1 through autoubiquitination and subsequent degradation.

In Hrd1, the Ser222-Asp258 loop is likely in close proximity to the proposed retrotranslocation channel^19,23^. Therefore, we tested whether ubiquitination of this cytosolic loop could also directly inhibit Hrd1 function by sterically obstructing substrate passage through the membrane. First, we confirmed that lysines within Ser222-Asp258 could be ubiquitinated in vitro. We purified wild-type Hrd1 and a destabilizing lysine substitution (Hrd1(N237K)), performed in vitro autoubiquitination, and found that Hrd1(WT) and Hrd1(N237K) both autoubiquitinated efficiently (Figure 5K). To determine whether Hrd1(N237K) was able to ubiquitinate residue 237, we excised polyubiquitinated Hrd1 and identified sites of lysine ubiquitination using mass spectrometry. In wild-type Hrd1, we identified ubiquitination sites near the cytosolic side of transmembrane segment 7 at Lys267 and Lys272 (Figure 5L). With Hrd1(N237K), we also observed ubiquitination at Lys267 and Lys272, but importantly observed two additional sites at the introduced N237K site and at Lys282 (Figure 5L). This directly supported the idea that Hrd1 can autoubiquitinate lysines within the cytosolic loop between transmembrane 6 and 7.

To test whether ubiquitination within the disordered cytosolic regions sterically inhibits Hrd1-mediated retrotranslocation, we used our in vitro retrotranslocation assay with Hrd1 containing lysine substitution within each disordered region (Hrd1(N237K-T435K)). We found that Hrd1(N237K-T435K) efficiently autoubiquitinated (Figures 5M and 5N), ubiquitinated externally-oriented substrates (Figures 5O and 5P), and retrotranslocated internally-oriented substrates, similarly to wild-type Hrd1 (Figures 5Q and 5R). These data demonstrate that the ubiquitination of these disordered cytosolic regions does not directly prevent substrate retrotranslocation. Rather, we concluded that the exclusion of lysines and cysteines from the transmembrane 6-7 loop and the disordered C-terminal region prevents premature degradation of Hrd1 catalyzed by autoubiquitination.

### The cytosolic loop between transmembrane segments 6 to 7 has a unique role in ERAD

Our work has uncovered clear similarities between the disordered C-terminal cytosolic region of Hrd1 and the disordered cytosolic transmembrane 6-7 loop. Furthermore, we defined an important role of the disordered C-terminal region in substrate interaction and retrotranslocation. As such, we hypothesized that the transmembrane 6-7 loop would also have a direct role in ERAD-L retrotranslocation and degradation. We returned to our deep mutational scanning data and found mutations near transmembrane 7 that were highly enriched but had high false discovery rates (Figures S2A and S2B). Nevertheless, we tested these variants against ERAD-L and ERAD-M substrates and found mutations that specifically inhibited ERAD-L substrate degradation (F268K, F268R, M269P, and I274T) (Figures 6A and S6A) without altered protein stability (Figure S6B).

**Figure 6.**
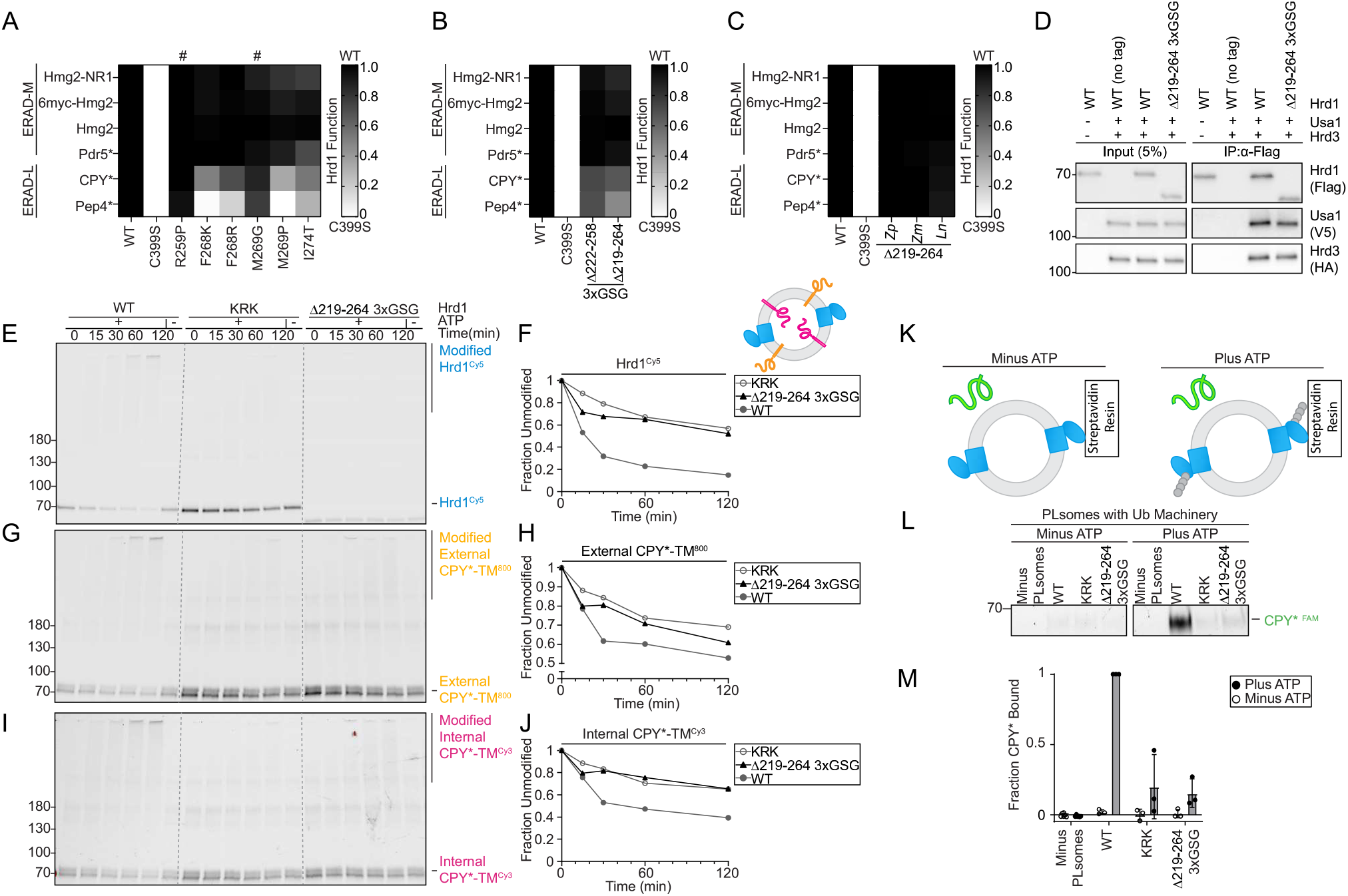
Cytosolic loop between transmembrane segments 6 and 7 required for ERAD-L degradation. A) Degradation of ERAD substrates by individual Hrd1 variants were followed by flow cytometry and summarized in a heatmap. Hrd1 variants were integrated in a *hrd1Δ* expressing individual ERAD substrates and subjected to a 4-hour mid-log chase. Wild-type Hrd1(WT) is set to 1 (full function, black) and inactive Hrd1(C399S) is set to 0 (no function, white). Values outside of this range were set to 0 or 1 (see Figure S6A). “#” indicate false positive hits. B) As in (B), but with the indicated Hrd1 regions replaced by a 3xGSG (Gly-Ser-Gly-Gly-Ser-Gly-Gly-Ser-Gly) linker sequence. C) As in (B), but with the indicated Hrd1 chimeras. Indicated region of *S. cerevisiae* Hrd1 was replaced with homologous region of *Zygosaccharomyces parabailii* (*Zp*), *Zygotorulaspora mrakii* (*Zm*), or *Lachancea nothofagi* (*Ln*). (See Figures S6C and S6D). D) Co-immunoprecipitation of the Hrd1 complex was performed with the indicated Hrd1 variants. 3xHA-Hrd3, 3xV5-Usa1, and Hrd1-3xFlag were integrated into *hrd1Δhrd3Δusa1Δ* cells, lysed, and immunoprecipitated with anti-Flag antibodies. Input represents 5% of the cleared lysate. This immunoblot is representative of 3 independent replicates. E) In vitro autoubiquitination of Hrd1^Cy5^ (blue) in a reconstituted proteoliposome system with externally-oriented CPY*-TM^800^ (orange), internally-oriented CPY*-TM^Cy3^ (magenta). Wild-type Hrd1 (WT-positive control), a retrotranslocation-defective Hrd1 (Hrd1(KRK)-negative control), or Hrd1(Δ219-264_3xGSG) were reconstituted and incubated with recombinant ubiquitination machinery for the indicated times. Samples were analyzed by SDS-PAGE and in-gel fluorescence scanning to visualize Hrd1. Red pixels indicate saturation of signal during the imaging. For second and third replicate see (Figures S6E-S6P). F) Quantification of E, showing unmodified Hrd1^Cy^^5^. G) As in E showing external CPY*-TM^800^. H) Quantification of (G), showing unmodified external CPY*-TM^800^. I) As in E showing internal CPY*-TM^Cy^^3^. J) Quantification (I), showing unmodified internal CPY*-TM^Cy^^3^. K) Schematic for soluble CPY* (green) interaction assay in proteoliposomes. Hrd1 is shown in blue and ubiquitin chains in gray. L) Hrd1 proteoliposomes (Plsomes) were immobilized at 1 µM and incubated with recombinant ubiquitination machinery, +/− ATP, washed, and incubated with 100nM CPY*^FAM^(green) (see K). Samples were eluted with biotin and analyzed by SDS-PAGE and in-gel fluorescence scanning. Red pixels indicate image saturation. M) Quantification of (L) displayed as mean +/− SEM, from three independent experiments. For this figure, the number of quantified replicates and individual values are shown in Table S8.

As with the C-terminal disordered region (Figure 3), we generated deletions of two regions (either Ser222-Asp258 or Gln219-Leu264), but only partially replaced the region with a 3xGSG linker (Gly-Ser-Gly-Gly-Ser-Gly-Gly-Ser-Gly). Hrd1(Δ222-258_3xGSG) degraded ERAD-M substrates normally but showed modest defects in ERAD-L substrate degradation (Figures 6B). The slightly expanded deletion (Hrd1(Δ219-264_3xGSG)) also degraded ERAD-M substrates normally but had stronger defects in ERAD-L substrate degradation (Figures 6B). When Gln219-Leu264 was replaced with homologous regions from diverging yeast strains (Figure S6D), we found that each chimera tested maintained a wildtype-like ability to degrade both ERAD-L and ERAD-M substrates (Figure 6C and S6C), even though many of these sequences were significantly shorter (Figure S6D).

We wondered why Hrd1(Δ219-264_3xGSG) failed to degrade ERAD-L substrates. First, we confirmed that Hrd1(Δ219-264_3xGSG) interacted normally with Usa1 and Hrd3 (Figure 6D) and had comparable protein levels to wild-type Hrd1 (Figure S6B). Then, we tested Hrd1(Δ219-264_3xGSG) using our in vitro retrotranslocation assay and observed severe defects because Hrd1(Δ219-264_3xGSG) was unable to autoubiquitinate, mirroring the negative control (Hrd1(KRK), Figures 6E and 6F). In addition, this variant barely ubiquitinated the externally-oriented substrate (Figures 6G and 6H) and failed to retrotranslocate internally-oriented substrate (Figures 6I and 6J). Because this variant could not autoubiquitinate efficiently, we expected that it would be unable to activate and expose the cytosolic substrate binding site that is in part formed by the C-terminal disordered region (Figure 4). To test this, we used the in vitro CPY* binding assay (Figure 6K) and found that Hrd1(Δ219-264_3xGSG) failed to interact with soluble CPY*, similarly to Hrd1(KRK) (Figures 6L and 6M). Taken together, in contrast to the disordered C-terminal region, the cytosolic transmembrane 6-7 loop was required for efficient Hrd1 autoubiquitination and therefore, each of the subsequent steps in Hrd1 retrotranslocation.

## Discussion

In this study, we developed a deep mutational scanning platform to identify critical residues for function of the Hrd1 ubiquitin ligase. This powerful approach allowed us to identify different clusters of mutations that disrupted Hrd1 function through distinct mechanisms (Figure 1). We identified mutations across the first two transmembrane segments that likely disrupt Hrd1/Hrd3 interaction, Hrd1/Der1 interaction, and by altering substrate specificity of the complex (Figure 2). We refined the Hrd1/Usa1 interaction site and identified a disordered C-terminal region of Hrd1 that interacts with ERAD-L substrates to facilitate retrotranslocation (Figure 3 and 4). Similarities to the newly identified C-terminal region directed us to another disordered cytosolic region between transmembrane segments 6 to 7 that is required for Hrd1 function (Figures 5 and 6). Degradation of ERAD-L substrates, but not ERAD-M substrates, requires both cytosolic disordered regions in vivo, but for different reasons. The disordered cytosolic loop between transmembrane segments 6 to 7 (Gln219-Leu264) promotes Hrd1 autoubiquitination, which is required for retrotranslocation. Following autoubiquitination, the disordered C-terminal region (Gly408-Thr480) promotes retrotranslocation by interacting with the substrate on the cytosolic side of the membrane. Substrate ubiquitination occurs in close proximity to these two regions, explaining why lysine and cysteine substitution are not tolerated there.

Based on our experiments, and previously published data^21–23^, we propose the following model (Figure 7). The Hrd1 complex engages substrate on the lumenal side of the membrane. The cytosolic loop between transmembrane segments 6 to 7 coordinates autoubiquitination to activate Hrd1 and open the cytosolic high-affinity substrate binding site(s), providing molecular interactions to enforce directionality and facilitate retrotranslocation, prior to substrate ubiquitination. These high-affinity sites ultimately position substrates for ubiquitination, allowing recruitment of the Cdc48 complex to improve the efficiency of retrotranslocation, extraction, and, ultimately, substrate degradation.

**Figure 7.**
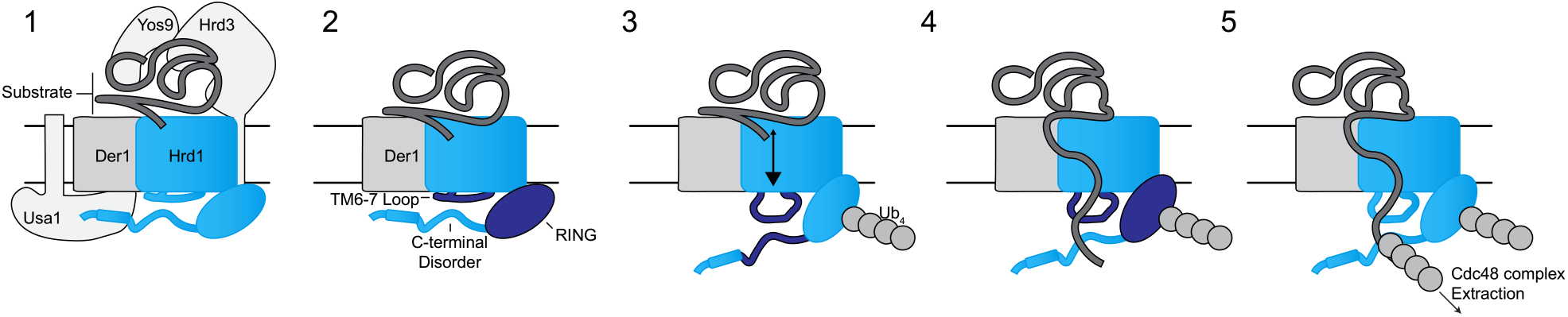
Model for retrotranslocation. Schematic for Hrd1 complex retrotranslocation. In stage 1 the Hrd1 complex recruits a lumenal substrate. In stage 2, lumenal substrate engages Der1-Hrd1 on the lumenal side of the membrane and Hrd1 autoubiquitination is driven by the cytosolic transmembrane 6-7 loop. In stage 3, Hrd1 autoubiquitination exposes the cytosolic substrate-binding sites in the transmembrane 6-7 loop and disordered C-terminal region creating a binding site for substrates on the cytosolic side of the membrane. In stage 4, Hrd1 ubiquitinates substrate and in stage 5 ubiquitinated substrate is extracted by the Cdc48 complex for degradation by the proteasome.

Initially, we were surprised to find new mutations in the cytosolic regions of Hrd1 to be important for retrotranslocation that were independent of Usa1 interaction or RING-finger domain formation because Hrd1 transmembrane segments can directly recognize substrate^18^. Furthermore, structural and biochemical studies have illuminated a proposed retrotranslocon channel formed by the transmembrane segments 3 through 8^19,21–23^. The mutations we found in transmembrane segments that specifically disrupted ERAD-L primarily fell within the first two transmembrane segments. The simplest explanation of these results is that mutations within transmembrane segment 2 would likely disrupt Hrd1/Der1 interaction. Transmembrane segment 1 mutations appear to alter the specificity of the complex, possibly by affecting the entry of substrate into the retrotranslocation channel^19^. Within the transmembrane segments that form the proposed retrotranslocation channel (TM3-8), we found relatively few mutations that specifically inhibited lumenal substrate degradation. We suspect that mutations within the channel itself affect both ERAD-L and ERAD-M substrate degradation or destabilize the Hrd1^17,42^. It is also likely that our method, focused on point mutations, would miss long-range or paired mutations with stronger phenotypes.

The cytosolic regions we have identified as essential for degradation of lumenal substrates are missing from current structural models of Hrd1^19,23^. In addition, innovative studies focused on the cytosolic domains of Hrd1 identified highly conserved structured regions that were essential for partner protein interactions, but the role of surrounding disordered regions themselves remained enigmatic^43^. While predicted disordered cytosolic regions are common to all forms of Hrd1, from fungi to humans (Figure S7A), their amino acid sequences have little conservation. This general trend was observed in our deep mutational scanning because many individual substitutions were tolerated in these regions, except for lysine and cysteine substitution. The apparent lack of sequence conservation and mutational tolerance, while maintaining ERAD function, is another reason why these regions may have gone undetected in previous studies. However, there are other ubiquitin ligases that use disordered domains to recognize their substrates^44^. Even in ERAD-related proteins, disordered domains appear to interact with integral membrane substrates^45^. Moreover, the existence of cytosolic substrate-interaction domains within Hrd1 was previously suggested, but the identity of this substrate interaction domain was unclear^22^. We identified a region in the C-terminal region of Hrd1 (Gly408-Thr480) that serves as this cytosolic ubiquitin-activated substrate-binding domain. Disrupting Gly408-Thr480 did not completely inhibit Hrd1’s ability to bind cytosolic substrate, consistent with the idea that another portion of Hrd1 (or even polyubiquitin chains themselves) could contribute additional affinity^22^. However, we propose that the cytosolic loop between transmembrane segments 6 to 7 (Gln219-Leu264) is another substrate interaction site because of its similar character to Gly408-Thr480 and the positioning directly adjacent to the retrotranslocation channel.

While our model is the simplest explanation of our results, more complicated models could also be applied to our findings. The Hrd1 complex is increasingly viewed as a dynamic complex with the precise active stoichiometry (or range of active stoichiometries) yet to be determined. Biochemical and structural data have demonstrated that Hrd1 can exist as monomers or dimers in complex with Der1. Moreover, activation of Hrd1 by ubiquitination may drive complex rearrangement^30^. The disordered domains of Hrd1 may be required for stabilizing transient complex formations or promoting the active Hrd1 stoichiometry.

Here, using deep mutational scanning combined with in vivo assays and in vitro reconstitution, we have demonstrated that *S. cerevisiae* Hrd1 requires the disordered cytosolic regions for function. It is important to note that all Hrd1 homologs, including mammalian Hrd1, have disordered cytosolic regions of varying lengths and positioning (Figure S7). We propose that these disordered cytosolic regions are broadly required for ER-associated degradation in all eukaryotes, but future studies will be needed to test this idea.

## Methods

### Strains and plasmids

Yeast strains used in this study were purchased from Horizon Discovery Ltd. and are derivatives of BY4741 (*MATa his3*Δ1 *leu2*Δ0 *met15*Δ0 *ura3*Δ0) or BY4742 (*MATα his3*Δ1 *leu2*Δ0 *lys2*Δ0 *ura3*Δ0) (Table S11). Yeast were cultured in synthetic dropout media (0.17 % (w/v) yeast nitrogen base(Becton, Dickinson and Company), 0.5 % (w/v) ammonium sulfate (Fisher), ∼0.1 % (w/v) drop-out powder (Teknova and Sigma), 2 % (w/v) glucose (Sigma)) at 30°C. Drop-out powder for synthetic complete media are at the following concentration: adenine sulfate (20 mg/L), uracil (20 mg/L), L-tryptophan (20 mg/L), L-histidine (20 mg/L), L-arginine (20 mg/L), L-methionine (20 mg/L), L-tyrosine (30 mg/L), L-leucine (60 mg/L), L-isoleucine (30 mg/L), L-lysine (30 mg/L), L-phenylalanine (50 mg/L), L-glutamic Acid (100 mg/L), L-aspartic Acid (100 mg/L), L-valine (150 mg/L), L-threonine (200 mg/L), and L-serine (400 mg/L). Combinational knockout strains were derived from transforming a PCR-amplified antibiotic targeting cassette with the LiAc/PEG methods or by crossing and sporulation^46^ and appropriate targeting was verified by PCR. Plasmids were constructed using either standard restriction cloning or HiFi DNA assembly (New England Biolabs) and propagated in DH5α *E. coli*. For most in vivo experiments, we used custom integrating cassettes targeted to the *leu2*Δ0, *his3*Δ1, or *ura3*Δ0 loci in BY4741/BY4742 strains. Centromeric plasmids were used only where specified in figure legends^47^. Note Pdr5* constructs used in Figure 2F, 3E, and 6C contained a second mutation A1174V that arose spontaneously during restriction cloning between vectors. All strains and plasmids available upon request (Tables S11 and S12).

### Hrd1 steady state collection

*hrd1Δ* cells expressing Hrd1-3xFlag (or the indicated Hrd1 variants) from the Hrd1 native promoter were grown with shaking at 30°C to mid-log phase between an OD_600_ of 0.35-1.05. Cells were pelleted, had 0.1 mm glass beads (BioSpec) added, frozen on dry ice, and stored at −80°C until lysis. Samples were resuspended in lysis buffer (10 mM 3-(N-morpholino)propanesulfonic acid (MOPS), pH 6.8, 1 % sodium dodecyl sulfate (SDS), 8 M urea, 10 mM ethylenediaminetetraacetic acid (EDTA), fresh protease inhibitors (1 mM Phenylmethylsulfonyl fluoride (PMSF), 1.5 µM pepstatin A)) at 25 OD_600_/mL. Samples were vortexed for 2 minutes then diluted with an equal volume of urea sample buffer (125mM trisaminomethane (Tris), pH 6.8, 4 % SDS, 8 M urea, 10 % β-mercaptoethanol). Samples were incubated at 65°C for 5 minutes before separating on SDS-polyacrylamide gel electrophoresis (SDS-PAGE), transferred to a polyvinylidene difluoride (PVDF) membrane, immunoblotted with anti-DYKDDDK (Genscript) and Mouse IgG HRP-linked whole Ab (Cytiva), then imaged using (ECL Select, Cytiva) with a ChemiDoc MP (Bio-Rad). For gel quantification, band intensities were measured using ImageJ (NIH), normalized to total protein in the sample detected using Bio-Rad Stain Free Dye Imaging Technology (StainFree). Hrd1 variant expression levels were normalized to wild-type Hrd1. All individual Hrd1 protein level values can be found in (Table S9). For statistical analysis one-way ANOVA tests were performed and p values were derived using Dunnett’s multiple comparisons test against Hrd1(WT) or a Welch’s t-test.

### Co-immunoprecipitation of Hrd1 Complex

Hrd1-3xFlag (or Hrd1 variants) were integrated at the *his3* locus, 3xHA-Hrd3 at the *leu2* locus, and 3xV5-Usa1 at the *ura3* locus in *hrd1Δhrd3Δusa1Δ* cells. Cells were cultured to mid-log phase, pelleted, resuspended in IP buffer (50 mM 2-[4-(2-Hydroxyethyl)-1-piperazine] ethanesulfonic acid (HEPES), pH 7.4, 150 mM potassium chloride (KCl)) with protease inhibitors (1 mM PMSF, 1.5 µM pepstatin

A) and flash frozen in liquid nitrogen to form yeast ‘balls’ prior to cryogenic lysis using freezer/mill (SPEX SamplePrep). Cell powders were thawed on ice and centrifuged at 18,000 x g for 10 min to collect the microsomal fraction. The pellets (P_18K_) were solubilized in IP buffer supplemented with 1 % decyl maltose neopentyl glycol (DMNG) and protease inhibitors by rotating for 1 hour at 4°C. Following solubilization, we collected the “input” samples, and 10 OD_600_ of solubilized proteins (180-200 µg proteins) were diluted 1:5 in IP buffer to a final concentration of 0.2 % DMNG. The solubilized proteins were mixed with 20 µL (40 µL slurry) anti-Flag M2 magnetic beads (Sigma) and rolled for 3 hours at 4°C. The bound proteins were washed 6-7 times with 1x volume of IP buffer (50 mM HEPES, pH 7.4, 150 mM KCl, 0.2 % DMNG) and eluted with 2x SDS-PAGE sample buffer. The samples were analyzed by SDS-PAGE and immunoblotting with anti-Flag (anti-DYKDDDDK, Genscript), anti-HA (clone 3F10, Roche), and anti-V5 (A01724, Genscript) antibodies with the inputs loaded at 5 %.

### Flow cytometry-based degradation assays

*hrd1*Δ cells expressing ERAD substrates were complemented with Hrd1, or Hrd1 variants. Cells were picked from transformation plates and subcultured in synthetic dropout media in 96 deep-well plates until cells entered log-phase (<1.5 OD_600_/ml). Cells were pelleted and resuspended in fresh synthetic dropout media supplemented with either cycloheximide (at 50 µg/ml), zaragozic acid (at 10 µg/ml, for Hmg2), or ethanol (0.1 % as a vehicle control) for the indicated times. At end of the time course, cells were pelleted, washed with ice-cold phosphate-buffered saline (PBS; 137 mM NaCl, 2.7 mM KCl, 10 mM sodium phosphate buffer, and 1.8 mM KH_2_PO_4_ (pH 7.4)), resuspended in PBS containing 1 µM SytoxBlue (viability dye; Invitrogen), and placed at 4°C during acquisition on either a Ze5 (Everest software; Bio-Rad) or a MACSQuant VYB (MACSQuantify software; Miltenyi Biotec). On the Ze5, the 488 nm laser was used for forward/side scatter to identify single cells, and Sytox Blue fluorescence was used to exclude dead cells from the 405 nm laser with a 460nm/22nm bandpass filter. GFP fluorescence was measured from the 488 nm laser with a 509nm/24nm bandpass filter, and mScarlet-I fluorescence was measured from the 561 nm laser with a 615nm/24nm bandpass filter. On the MacsQuant VYB we used the 561 nm laser for forward/side scatter to identify single cells, and SytoxBlue fluorescence was used to exclude dead cells from the 405 nm laser with a 450nm/50nm bandpass filter set. GFP fluorescence was measured from the 488 nm laser with a 525nm/50nm bandpass filter set, and mScarlet-I fluorescence from the 561 nm laser with a 615nm/20nm bandpass filter set. FlowJo V10.7.1 (FlowJo LLC) was used to analyze FCS files (version 3.1) on FSC/SSC to eliminate debris, gate on single cells, and eliminate dead cells. median GFP, or mScarlet-I, fluorescence values from at least 10,000 cells were exported and used to quantify Hrd1 variant function. For quantification, we determined the fraction of substrate remaining after the treatment, compared to the vehicle control. The fraction of substrate remaining was normalized to wild-type Hrd1 set to “1”, and Hrd1(C399S) set to “0”. All individual flow cytometry functional values can be found in (Table S8). For statistical analysis one-way ANOVA tests were performed and p values were derived using Dunnett’s multiple comparisons test against Hrd1(WT).

For saturated chases, single cells were inoculated in synthetic dropout media and grown overnight (∼14 hours). Cells were diluted 1 to 50, or 1 to 100, in fresh synthetic dropout media and cultured for ∼24 hours to enter a “saturated chase” and processed as described above.

Flow cytometry plots were generated using FlowJo V10.7.1, and all flow cytometry data are from at least 10,000 events passing FSC/SSC and viability gates.

### Library generation

Tiling primers mutagenesis was performed as previously described^37^. We mutagenized Hrd1 in five regions (amino acids: 1-110, 111-220, 221-330, 331-440, 441-551), to enable 2 x 300bp Illumina sequencing across the regions. A linearized PCR template was generated by digesting a wild-type Hrd1 plasmid with restriction enzyme and purified using a QIAquick PCR purification kit (Qiagen). For each region two separate PCR reactions were set up, one for forward tiling primers and one for reverse tiling primers amplifying the linearized wild-type Hrd1 sequence with a flanking primer adding homology for gap-plasmid repair in yeast (Table S13 and S14). For library generation, the first round of PCR consisted of 7 cycles of the following program using Q5 High-Fidelity DNA Polymerase (New England Biolabs). Step1: 98°C for 2 min; step 2: 98°C for 30 sec; step 3: 72°C for 1 sec; step 4: 60°C for 30 sec, cooling at 0.5°C/sec; step 5: 72°C for 30 sec; step 6: 7 cycles of return to step 2; step 7: 72°C for 1 min.

These PCR products were diluted 1 to 4 with water, and each forward and reverse library was combined as the template for a second round of PCR using the flanking primers. Step1: 98°C for 2 min; step 2: 98°C for 30 sec; step 3: 72°C for 1 sec; step 4: 60°C for 30 sec, cooling at 0.5°C/sec; step 5: 72°C for 30 sec; step 6: 20 cycles of return to step 2; step 7: 72°C for 1 min. The amplified libraries were cleaned up by a QIAquick PCR purification kit.

For the vector to be used for recombination, centromeric plasmids containing the native promoter and terminator of Hrd1 had partial replacement of coding sequence corresponding to amino acids 1-110, 111-220, 221-330, 331-440, or 441-551 with an EcoRI restriction site. Plasmids were linearized with EcoRI-HF and purified using QIAquick PCR purification kit. This method produced a range of mutations per region (Table S10).

To generate the mutagenized libraries yeast cells expressing integrated substrates were transformed with a 1:5 molar ratio of linearized vector to PCR fragments product using standard LiAc/PEG transformation using homologous recombination for gap-plasmid repair strategy^38^. Post transformation, 0.1 % of the cell mixture was plated on synthetic dropout plates to determine transformation efficiency and the rest was inoculated into synthetic dropout media and cultured for two days to reach saturation (Table S1).

### Fluorescence-activated cell sorting (FACS)

Two days after the library generation, the cell libraries were diluted 1:100 in fresh synthetic dropout media and grown for 24 hours, allowing cells to enter into a “saturated chase”. Prior to FACS, we collected an aliquot of cells as the “input” library, and stored the cells at −80°C. For FACS, cells were kept at room temperature, pelleted, washed in PBS, resuspended in PBS containing 1µM SytoxBlue, and sorted using a Bigfoot Spectral Cell Sorter (Invitrogen (formally Propel Labs)). Forward and side measurements were analyzed on the 488nm laser to gate for single cells and SytoxBlue fluorescence was used to exclude dead cells from the 445nm laser with a 465nm/22nm bandpass filter. GFP fluorescence was measured from the 488nm laser with a 507nm/19nm bandpass filter set and mScarlet-I fluorescence was measured from the 561nm laser with a 605nm/15nm bandpass filter set. Cells in the wildtype-like sort bin (WT) and ERAD-L defective sort bin (L) were collected in tubes containing 2x synthetic dropout media to aid cell recovery. Over 3.3 million cells passing FSC/SSC and viability gating were sorted over (see Table S1 for the number of cells sorted into each bin).

Sorted cells were grown in synthetic dropout media to saturation and an aliquot of cells was collected and stored at −80°C prior to genomic extraction. This frozen aliquot of cells represents the sorted population used for NGS library prep and analysis (wildtype-like or ERAD-L defective populations). The remaining cells were subjected to a saturated chase to confirm phenotype the next day. Some ERAD-L defective populations were subjected to a second round of sorting to enrich for the lumenal defective phenotype (Table S1).

### Illumina sequencing and data analysis

DNA was extracted from 30-40 million (3-4 OD_600_ equivalents) cells using a zymolyase method^48^. Briefly, cells were resuspended in 100 µL of buffer Z (50 mM Tris, pH 7.4 and 1-3 mg/ml of zymolyase 100T (Amsbio)) and shaken at 37°C for 1-2 hours. The solution was heated at 95°C for 6 minutes before centrifugation to clear insoluble material.

NGS libraries were prepared for amplicon sequencing on an Illumina platform. Libraries were generated from two PCR reactions both using Q5 High-Fidelity DNA Polymerase (NEB). Primers flanking mutated regions amplified, 25 cycles, the extracted DNA from 3-4 million cells (10 µL of extraction solution) while adding on partial R1 and R2 read adaptors (Table S14). The second round of PCR (8-9 cycles) added i5 and i7 indexes using the Index Kit 2 for Illumina (Apexbio Technology LLC). The libraries were cleaned up using AMPure XP beads, normalized, pooled, and submitted for sequencing on a MiSeq 2×300 platform through Genewiz (now Azenta Life Sciences).

For analysis, de-multiplexed FASTQ files were provided from Azenta. These files were trimmed with cutadapt (using the following arguments: -a CTGTCTCTTATACACATCT -A CTGTCTCTTATACACATCT -u 8 -U 8 -q 30), 3’ overlapping pair-end reads were merged with fastp (using the default settings), and aligned to a wild-type Hrd1 sequence with bowtie2 (using the “very-sensitive” setting), and resulting SAM files was sorted on name while converting to a BAM file using samtools^49–52^. The sorted BAM files were used as input for python scripts that translated and counted Hrd1 mutations for all sequences without insertions or deletions (indels) identified during Bowtie alignment.

To test variant enrichment in the ERAD-L defective Hrd1 sort bin, we only analyzed reads with single amino acid substitutions. We performed rate ratio tests using the test_poisson_2indep function from version 0.13.5 of the python module statsmodels.stats.rates. Arguments to the test_poisson_2indep function were as follows: “count1”: the number of reads in the ERAD-L defective phenotype with amino acid ‘a’ at Hrd1 position ‘p’, “exposure1”: the total number of reads arising from the ERAD-L defective phenotype, “count2”: the number of reads in the input mutant library with amino acid ‘a’ at Hrd1 position ‘p’, “exposure2”: the total number of reads in the mutant library, “alternative”: “greater”. We used jackknife sampling to incorporate unpaired replicates, performing a separate set of hypothesis tests for each jackknife replicate to arrive at our final estimates of p-values and variant enrichment in the ERAD-L defective bin. We used the multipletests function in version 0.13.5 of the python module statsmodels.stats.multitest to correct for multiple hypothesis testing using the method of Benjamini and Hochberg^53^. Variant enrichment (or depletion) in the wildtype-like bin was analyzed similarly as the ERAD-L defective bin, but the “alternative”: was set to “two-sided”.

### Screening optimization mutant isolation

Hrd1(F46R) was isolated with a similar method as the DMS screen above with the following differences. Hrd1 was mutagenized using tiling primers from across the first 384 amino acids and transformed into *hrd1Δ* expressing substrates from centromeric plasmids. Cell sorting occurred using a MoFlo Astrios (Beckman). Forward and side measurements were analyzed on the 488nm laser to gate for single cells. GFP fluorescence was measured from the 488nm laser with a 513nm/26nm bandpass filter set and mScarlet-I fluorescence was measured from the 561nm laser with a 614nm/20nm bandpass filter set. Sorted cells were plated and individual colonies were confirmed for ERAD-L defective phenotype before Hrd1 plasmid isolation (Zymoprep Yeast Plasmid Miniprep II) and Sanger-sequencing.

Hrd1(T416P,W481L,I505T) was isolated as follows. Hrd1 was mutagenized across the entire coding sequence and cloned into a centromeric plasmid using HiFi Assembly (New England BioLabs). The centromeric Hrd1 library was transformed into *hrd1Δ* expressing substrates from centromeric plasmids and plated. Plates were fluorescently imaged using a ChemiDoc MP (Bio-Rad). Individual colonies appearing ERAD-L defective were selected and confirmed using flow cytometry before Hrd1 plasmid isolation (Zymoprep Yeast Plasmid Miniprep II) and Sanger-sequencing.

### Strains and plasmids for protein expression

Uba1 was purified from the InvSc1 strain (Invitrogen). Both CPY*-TM and soluble CPY* were purified from a *hrd3*Δ*alg3*Δ strain derived from BY4741 (yRB129: *MAT*a *his3*Δ1 *leu2*Δ0 *ura3*Δ0 *met15*Δ0 *hrd3Δ::kanRMX4 alg3Δ::hphNT1*). Hrd1 was expressed and purified from a *hrd1*Δ*ubc7*Δ diploid (yBGP55B: *MAT*a/α *his3*Δ1/*his3*Δ1 *leu2*Δ0/*leu2*Δ0 *LYS2*/*lys2*Δ0 *met15*Δ0/*MET15 ura3*Δ0/*ura3*Δ0 *hrd1*::*HphNT1*/*hrd1*::*HphNT1 ubc7*::*KanRMX4*/*ubc7*:*KanRMX4*). Bacterial expression strains were BL21-CodonPlus (DE3) RIPL (Agilent). (See Table S11)

Yeast overexpression for purification were from 2μ plasmids of the pRS42X series driven by the inducible Gal1 promoter^54^. Bacterial expression of Cue1 and Ubc7 were expressed from two different fusion proteins. As previously described^18^, where Cue1(24-203) (pAS153) and Ubc7 (pAS159) were expressed with an N-terminal His_6_ tag in a pET28B vector (Novagen). Cue1 and Ubc7 were also expressed from a K27-His_14_-pSUMO vector allowing for the complete removal of affinity and solubility tags^55^. (See Table S12)

### Yeast protein expression and purification

Yeast strains with plasmids for the expression of proteins of interest were grown at 30°C with shaking in synthetic dropout media. 1:150 dilutions of actively growing cultures were used to inoculate eight 1L cultures in 2.8L Fernbach flasks. The cultures were grown for 24 hours at 30°C with shaking. To induce expression, 4x yeast extract peptone (YP) broth containing 8 % (w/v) galactose was added to the culture to a final concentration of 1x YP and 2 % galactose. The temperature was shifted to 25°C, and the culture was grown for 15 to 18 hours. Cells were pelleted, washed with water or 2 mM dithiothreitol (DTT) (to weaken the cell wall), flash-frozen with liquid nitrogen, and stored at −80°C.

All purification steps were conducted at 4°C unless otherwise specified. Approximately 150 to 200 grams of yeast cells were resuspended in buffer Y_1_ (50 mM HEPES, pH 7.4, 300 mM KCl, and 0.5 mM tris(2-carboxyethyl)phosphine (TCEP)) with freshly supplemented protease inhibitors (1 mM PMSF, 1.5 µM Pepstatin A, 0.2 mM diisopropyl fluorophosphate (DFP)). 0.5 mm glass beads were added to ⅓ to ½ the volume of the cell suspension and were subjected to bead beating (BioSpec), either split across two bead beater rounds or unbroken cells from the first bead beating round were subjected to a second round of bead beating. Cells were lysed by beat beating for 20 minutes of 20 seconds on/ 40 seconds off. The lysate was cleared with two low-speed spins (2,000 x g for 10 minutes), and the supernatant was centrifuged at 42,000 rpm for 33 minutes using a 45Ti rotor (Beckham, RCF average 138,000 x g). For Hrd1 purification, the membrane fraction was washed twice by resuspending the membrane fraction in 180 mL of buffer Y_1_ containing protease inhibitors (1 mM PMSF, 1.5 µM Pepstatin A, 0.1 mM DFP) and pelleting using 45Ti ultracentrifugation.

For Hrd1 purification, the membrane fraction was resuspended in buffer 180ml of Y_1_ with freshly supplemented protease inhibitors (1 mM PMSF, 1.5 μM Pepstatin A, 0.1 mM DFP) and 1 % (w/v) DMNG. The membrane was rolled for 1 hour, and insoluble material was removed by ultracentrifugation in a 45Ti (42,000 rpm for 33 minutes). The supernatant was incubated with affinity resin (1.5 ml of Streptavidin Agarose (Pierce)) and concurrently sortase-labeled overnight (∼12 to 15 hours), using the sortase A (P94R/D160N/D165A/K190E/K196T) pentamutant^56^ and Sulfo-Cy5-Maleimide (Lumiprobe) coupled to a Gly-Gly-Gly-Cys peptide (Genscript). The next morning, the streptavidin agarose was washed seven times with 50 mL of buffer Y_1_ containing 1 % (w/v) DMNG, then 1 mM DMNG, and the remaining washes with 120 µM DMNG. The fourth wash was conducted at room temperature with 0.5 mM adenosine triphosphate (ATP) supplemented. Hrd1 was eluted off streptavidin resin in Buffer Y_1_ supplemented with 120 µM DMNG and 2mM biotin. Elutions were pooled based on yield and purity, as assessed by SDS-PAGE and Coomassie blue staining. Pooled elutions were concentrated and further purified using size exclusion chromatography (Superose 6 Increase 10/300 GL (Cytiva Life Sciences)) with 25 (or 50) mM HEPES, pH 7.4, 300 mM KCl, 0.25 (or 0.5 mM) TCEP, and 120 µM DMNG. Peak fractions were pooled, concentrated, and flash-frozen in liquid nitrogen.

For soluble CPY* with C-terminal sortase recognition tag, 3C cleavage site, and His_14_ affinity tag, we collected membranes as described above. The membrane fraction was resuspended under denaturing conditions in buffer Y_2_ (50 mM HEPES, pH 7.4, 300 mM KCl, 30 mM imidazole, 0.5 mM TCEP, and 6M urea) with fresh protease inhibitors and rolled for 1 hour. The urea insoluble material was cleared by 45Ti ultracentrifugation (42,000 rpm, 33 minutes). The urea-soluble supernatant was rolled with 10 mL of HisPur Ni-NTA Resin (Thermo Scientific) for 1.5 hours. The nickel resin was washed ten times with 50 mL of buffer Y_3_ (50 mM HEPES, pH 7.4, 300 mM KCl, 0.5 mM TCEP, 30 mM imidazole) containing the following concentration of urea per wash: wash 1, 6M urea; wash 2-4, 3M urea; wash 5-8, 1M urea; and wash 9-10, no urea. The CPY* material was eluted with 400 mM imidazole under denaturing conditions in buffer Y_2_. The denatured CPY* was refolded over a 20-hour dialysis. First, the 6M urea was diluted linearly to 3M urea using buffer Y_1_ over a ∼6 hour period. Next, 3M urea was diluted linearly to 0.5 M urea over 10 hours. Finally, the 0.5M urea solution was dialyzed twice against buffer Y_1_ bringing the final urea concentration to ∼1mM. The CPY* was labeled overnight (∼14 hours) with FAM-maleimide (Lumiprobe) coupled to Gly-Gly-Gly-Cys peptide using Sortase A (P94R/D160N/D165A/K190E/K196T) that replaces the C-terminal 3C cleavage site and His_14_ affinity tag if labeled. The overnight sortase labeling caused about ⅓ of the CPY* to precipitate out of solution. Insoluble CPY*-FAM was collected by low-speed centrifugation and re-solubilized in buffer Y_2_ for 1 hour. The solubilized denatured CPY* was depleted of His_14_ tag or full-length His_14_ containing species by three successive passes over 2mL of HisPur Ni-NTA Resin (Thermo Scientific). The urea was dialyzed out over 20 hours bringing the final urea concentration to ∼1mM in buffer Y_1_, as described above. The protein was concentrated, aliquoted, and flash-frozen.

CPY*-TM was purified as described previously^21^. Briefly, the membrane fraction was resuspended in 200 ml of buffer R_1_ (50 mM HEPES, pH 7.4, 300 mM KCl, 1 mM MgCl_2_, 1 mM TCEP, 6 M urea, 1 % tridecylphosphocholine (Fos-choline 13, Fos13), 30 mM imidazole) for 60 minutes. Insoluble material was removed by 45Ti ultracentrifugation (42,000 rpm, 30 minutes). Soluble material was incubated with His60 Superflow resin (Clontech) for 60 minutes. The resin was washed with 10 column volumes (CV) of buffer R_2_ (25 mM HEPES, pH 7.4, 300 mM KCl, 1 mM MgCl_2_, 0.5 mM TCEP, 2 mM Fos13, 30 mM imidazole) supplemented with 6 M urea. The column was washed with decreasing amounts of urea: 10 CV of R_2_(5 M Urea), 10 CV of R_2_(3 M Urea), 10 CV of R_2_(1 M Urea), and 20 CV of R_2_(No Urea). The protein was eluted with 400 mM imidazole in buffer R_2_, labeled with Dylight800 via Sortase A, and purified by gel filtration.

For Uba1 with N-terminal His_14_ affinity and TEV cleavage site, cell lysis occurred as described above with the following modifications. First, cells were resuspended in buffer Y_4_ (50 mM HEPES, pH 7.4, 300 mM KCl, 30 mM imidazole) with fresh protease inhibitors (1 mM PMSF, 1.5 µM Pepstatin A, 1x protease inhibitor cocktail (PIC)). Second, the supernatant, not membrane fraction, was taken following ultracentrifugation. The supernatant was rolled with 7.5 ml of HisPur Ni-NTA Resin (Thermo Scientific). The resin was washed with 25 times the bed volume with buffer Y_4_. Uba1 was eluted with 400 mM imidazole in buffer Y_4_. 5mM DTT was immediately added to each elution. Elutions were pooled based on yield and purity, as assessed by SDS-PAGE and Coomassie blue staining. The pooled elutions were supplemented with 5% glycerol and the His_14_ tag was removed using TEV protease (1 to 100 molar ratio) overnight (∼14 hours). The next morning, cleaved Uba1 was purified by ion-exchange chromatography using a HiTrap MonoQ column (Cytiva). We used a linear gradient of 50 mM KCl to 1000 mM KCl in 50 mM HEPES, pH 7.4; the Uba1 eluted around 350 mM KCl. The peak fractions were pooled, concentrated, and separated by size-exclusion chromatography (HiLoad 16/600 Superdex 200 pg (GE Healthcare, now Cytiva) in 20 mM HEPES, pH 7.4, 150 mM KCl, 300 mM sorbitol, and 0.5 mM TCEP. Peak fractions were pooled, concentrated, aliquoted, and flash-frozen in liquid nitrogen.

### Bacterial Protein Expression and Purification

Ubc7 and Cue1(Gln24-Thr203) were expressed and purified in two different ways. The first method was as described previously^18^.

K27 His_14_-SUMO Ubc7 was expressed in *E. coli* BL21-CodonPlus (DE3) RIPL cells (Aligent). Cells were inoculated into starter cultures containing kanamycin (50 µg/ml) and chloramphenicol (25 µg/ml) at 37°C with shaking and grown overnight (∼14 hours). The cultures were diluted 1:150 into 2.8L Fernback flasks with 1 L of terrific broth (TB) containing kanamycin and chloramphenicol at 37°C. When the cells reached 0.8-1.0 OD_600_/ml, protein expression was induced with 0.5 mM Isopropyl β-D-1-thiogalactopyranoside (IPTG). The cultures were grown at 18°C and 220 rpm for 17.5 hours. Cells were collected, washed in water, frozen on dry ice, and stored at −80°C. K27 His_14_-SUMO Cue1(Gln24-Thr203) was expressed as above with the following adjustments. Chloramphenicol was at 35µg/ml, induction occurred at 0.55 OD_600_/ml, and cells were grown at 18°C for 18.5 hours.

For K27 His_14_-SUMO Ubc7, the cell pellet (97 grams) was thawed and resuspended in 180 mL of buffer B_1_ (50 mM Tris, pH 8, 500 mM NaCl, 0.25 mM TCEP, and 30 mM imidazole) with fresh protease inhibitors (1 mM PMSF, 1.5 μM Pepstatin, 1x PIC). Cells were lysed via sonication, and the cell lysate was subjected to ultracentrifugation using a 45Ti (42,000 rpm, 33 minutes). The supernatant was incubated with 7.5mL of HisPur Ni-NTA resin for 2.5 hours and washed six times with buffer B_1_. The resin was washed with an additional 75 mL of buffer B_2_ (25 mM HEPES pH 7.4, 300 mM KCl, and 0.25 mM TCEP) supplemented with 30 mM imidazole. Ubc7 was eluted in 7.5 mL batches with 400 mM imidazole in buffer B_2_. An additional 2mM of TCEP was added to each elution. The peak fractions were pooled, and Ulp1 (SUMO-protease) was added to 4 μM and incubated overnight to cleave the His_14_-SUMO tag, while dialyzing against buffer B_2_ (to remove imidazole). Following overnight incubation, precipitated Ubc7 was filtered from solution. The His_14_-SUMO tag was removed from the solution, by passing the solution over 5 mL of HisPur Ni-NTA resin three successive times. Ubc7 was further purified by ion exchange chromatography using a HiTrap monoQ HP column(Cytiva) with a salt gradient from 50 mM KCl to 530 mM KCl in 25 mM HEPES pH 7.4 containing 0.25 mM TCEP; Ubc7 eluted around 235 mM of KCl. Peak fractions were pooled, concentrated, aliquoted, and flash-frozen in liquid nitrogen.

The K27 His_14_-SUMO Cue1(Gln24-Thr203) cell pellet (12 grams) was resuspended in 180 mL of buffer B_3_ (50 mM HEPES, pH 7.4, 300 mM KCl, 6 M urea, 10 mM EDTA) with fresh protease inhibitors (1 mM PMSF, 0.1 mM DFP, 1.5 μM pepstatin, 0.5 μM Bestatin, 1x PIC). The cells were lysed using sonication, and the insoluble material was cleared via ultracentrifugation in a 45Ti (42,000 rpm 33 minutes). The supernatant was collected, supplemented with 5 mM imidazole, and incubated with 4mL cOmplete His-Tag Purification Resin (Roche) for 1 hour. The resin was washed with 75x bed volume with buffer B_3_ supplemented with 5mL imidazole and eluted in batches of 6 mL of B_3_ supplemented with 400 mM imidazole. 2 mM TCEP was immediately added to the elutions. Cue1 was refolded by removal of the urea using dialysis. 6 M urea was dialyzed linearly to 1.5 M urea over a period of 3 hours using buffer B_4_ (25 mM HEPES pH 7.4, 300 mM KCl, 0.25 mM TCEP). 1.5 M urea was dialyzed to ∼142 mM urea over 14.5 hours. The solution was dialyzed against buffer B**_4_** for 1 hour bringing the final urea to ∼4.5 mM. We observed no precipitation of His_14_-SUMO Cue1. Ulp1 was added at a 1:75 (mass:mass) and incubated for 1 hour. To remove the His_14_-SUMO tag, the solution passed successively over 1.5 mL of HisPur Ni-NTA resin three times. Cue1 was further purified by ion exchange chromatography using a HiTrap monoQ HP column (Cytiva) with a salt gradient from 20 mM to 520 mM in 25 mM HEPES, pH 7.4 with 0.25 mM TCEP; Cue1 eluted around 150 mM KCl. Peak elutions were pooled, concentrated, aliquoted, and flash-frozen in liquid nitrogen.

### In vitro ubiquitination

In vitro ubiquitination occurred at 30°C in 25 mM HEPES, pH 7.4, 150 mM KCl, 5 mM MgCl_2_, 0.1 mM TCEP, and 120 μM DMNG. The ubiquitination machinery was added at the following concentrations: 0.2 μM Uba1, 2 μM Ubc7, 2 μM Cue1(Gln24-Thr203), 0.5 μM Hrd1, 125 μM ubiquitin (yeast recombinant from bio-techne (R&D Systems)), and 0.6 μM bovine serum albumin (BSA; A3311 from Sigma). 2mM ATP was added to start the reaction. The samples were analyzed on SDS-PAGE and fluorescence scanning (ChemiDoc MP, Bio-Rad). The gel section corresponding to poly-ubiquitinated Hrd1 was excised and sent for ubiquitin site identification at the Taplin Mass Spectrometry Facility (Harvard Medical School).

Excised gel bands were reduced with 1 mM DTT for 30 minutes at 60°C, alkylated with 5 mM iodoacetamide for 15 minutes at room temperature, dehydrated with acetonitrile, and dried in a speed-vac. Gels pieces were rehydrated in 50 mM ammonium bicarbonate, and peptides were generated by in-gel digestion using trypsin (sequencing-grade (Promega)) at 37°C overnight. Peptides were extracted in 50 % acetonitrile and 1 % formic acid, the peptide solution was dried in a speed-vac for 1 hour and stored at 4°C until analysis.

On day of analysis, samples were reconstituted in 5-10 μL of HPLC solvent A (2.5 % acetonitrile, 0.1 % formic acid). Samples were separated on a nano-scale reverse-phase HPLC capillary column (100 μm inner diameter x ∼25 cm length) packed with 2.6 μm C18 spherical silica beads. Elution occurred over a gradient of increasing HPLC solvent B (97.5 % acetonitrile, 0.1 % formic acid). As peptides eluted off column, they were subjected to electrospray ionization and entered an LTQ Orbitrap Velos Pro ion-trap mass spectrometer (ThermoFischer). Peptides were detected, isolated, and fragmented to produce a tandem mass spectrum of specific fragment ions for each peptide. Peptide sequences are matched to protein sequences by Sequest (ThermoFinnigan). The modification of 114.0429 mass units to lysine was included in the database searches to determine ubiquitin-modified peptides. Data was filtered with a 1 % false discovery rate.

### In vitro retrotranslocation

1,2-dioleoyl-sn-glycero-3-phosphocholine (DOPC) from Avanti Polar Lipids was solubilized in chloroform and aliquoted. The solvent was removed under a nitrogen stream, followed by placing the aliquot into a lyophilizer overnight (∼14 hours). DOPC was resuspended at 6.2 mM in buffer R (25 mM HEPES, pH 7.4, 150 mM KCl, 1 mM MgCl2, 100 μM TCEP) by vortexing. Lipids were completely solubilized in 1.2% (w/v; 14.8mM) Triton X-100 (Anatrace) for 20 min at room temperature. CPY*-TM substrate labeled with Dylight800 was added to 2.5μM bringing DOPC to 5.1 mM, and the mixture was incubated at 4°C with gentle shaking for 60 min. The detergent was removed by three 2-hour incubations with ∼33 mg of Bio-Beads SM2 (Bio-Rad) followed by overnight incubation with ∼50 mg of Bio-Beads SM2 with all incubations occurring at 4°C.

With sealed liposomes, color switching internal-CPY* occurred by adding 10 mM CaCl2, 50 μM sulfo-cyanine3-maleimide (Cy3) (Lumiprobe) coupled to a Gly-Gly-Gly-Cys peptide, and 2 μM Sortase A for 1 hour at 4°C. The proteoliposomes were mixed 1:1 (vol:vol) with 80 % glycerol in buffer R. The proteoliposomes were overlaid with 30% glycerol, 15 % glycerol, 5 % glycerol, and 0 % glycerol (all prepared in buffer R). The step gradient was centrifuged for 3 hrs at 50,000 rpm in a TLS-55 rotor (RCF_avg_ 166,180). Five fractions were collected, starting from the top of the gradient. Proteoliposomes formed a sharp band between the 15 % and 5 % glycerol layers. Sortase A and GGGC-fluorophore stayed in the 40 % glycerol layer.

To incorporate Hrd1 into the proteoliposomes, the floated samples were partially solubilized by the addition of 0.1 % (w/v) DMNG on ice for 30 min. Hrd1 was added to 2 μM, and the mixture was incubated on ice for 60 min. DMNG was removed by five successive passes over detergent removal resin (Pierce) at a ratio of 2.75x resin to proteoliposome material. The proteoliposomes were then mixed 1:1 with 80 % glycerol in buffer R and subjected to a glycerol step gradient centrifugation consisting of 30 %,15 %, and 0 % glycerol layers (prepared in buffer R).

The proteoliposomes containing CPY*-TM and Hrd1 floated to an interface between 15 % and 0 % glycerol layers. The liposomes were diluted 1 to 2 (or 1 to 4) in buffer R. The liposome solution was brought to room temp for 5 minutes, and then liposomes were added to an equal volume of ubiquitination machinery at the following concentrations: 0.4 μM Uba1, 4 μM Ubc7, 4 μM Cue1(Gln24-Thr203), 100 μM ubiquitin, and 0.6 μM BSA. The retrotranslocation assay was initiated with the addition of 2 mM ATP, and reactions were incubated at 30°C. The samples were analyzed on SDS-PAGE and fluorescence scanning ChemiDoc MP (Bio-Rad). Quantification of band intensity occurred using ImageJ, and the fraction unmodified was calculated by taking band intensity at indicated time point over band intensity at the zero-minute time point.

### Hrd1 CPY* Binding

DOPC, as prepared for retrotranslocation assays, was resuspended in buffer C_1_ (25 mM HEPES, pH 7.4, 150 mM KCl, 5 mM MgCl, 0.1 mM TCEP). Liposomes were extruded through 100 nm polycarbonate membrane, partially solubilized with 0.1 % DMNG for 30 minutes on ice, and incubated with 2 μM Hrd1 for 1 hour on ice. DMNG was removed by four successive passes over detergent removal resin (Pierce) at a ratio of 3.35x resin to proteoliposome material. The proteoliposomes were then mixed 1:1 with 80 % glycerol prepared in buffer C_1_ and subjected to glycerol step gradient centrifugation consisting of 30 %, 15 %, 5 %, and 0 % glycerol layers (prepared in buffer C_1_) in a TLS-55 (50,000 rpm for 3 hours). The liposomes with incorporated Hrd1 floated to the interface between the 15 % and 5 % glycerol layers.

Pierce magnetic streptavidin beads were prewashed with 112.5 μL of buffer C_1_ supplemented with 2 mg/ml BSA and 2 mM DOPC, followed by 225 μL of buffer B_1_ per 7.5 μL of beads. 20 μL of 1 μM Hrd1 proteoliposomes were bound to 7.5 μL of prewashed beads at 30°C for 30 min. Background binding controls lacking proteoliposomes were incubated with buffer C_2_ (25 mM HEPES, pH 7.4, 150 mM potassium chloride, 5 mM magnesium chloride, 0.1 mM TCEP, 0.6 μM BSA). Unbound Hrd1 proteoliposomes were pipetted off, and immobilized Hrd1 proteoliposomes were resuspended in ubiquitin mix consisting 0.2 μM Uba1, 2 μM Ubc7, 2 μM Cue1(Gln24-Thr203), and 50 μM ubiquitin in buffer C_2_. Ubiquitination was initiated by the addition of 2 mM ATP and incubated at 30°C for 1hr. Beads were subsequently washed with 225 μL of buffer C_2_ to remove the ubiquitination mix. The substrate, 20 μL of 0.1μM CPY*-FAM, was incubated with immobilized Hrd1 proteoliposomes at 30°C for 1 hr. The unbound CPY*-FAM was collected, and the beads were washed with 225 μL of buffer C_2_. The bound material was eluted with 2 mM biotin in 50 mM MOPS for 5 minutes at 30°C. The bound samples were analyzed by SDS–PAGE and fluorescence scanning ChemiDoc MP (Bio-Rad). Substrate binding was quantified from three independent experiments using ImageJ. To determine the fraction bound, band intensity was normalized to CPY*-FAM bound to Hrd1(WT) with ATP condition.

## Acknowledgments

We would like to thank Devon Dennison and other members of the Baldridge Lab for critical reading of the manuscript and thoughtful discussion. We thank former lab member Morgan Glaser for cloning the Hrd1 libraries used to isolate Hrd1(T416P,W481L,I505T). We thank Dave Adams and other members of the Michigan Flow Cytometry Core for assistance with flow cytometry training and cell sorting. We also thank Ross Tomaino at the Harvard Taplin Mass Spectrometry Facility for assistance in Hrd1 ubiquitination site identification.

## Author Contributions

B.G.P. conceptualized the project, developed the methodology, performed the investigation, analyzed the data, visualized the data, wrote the original draft, and reviewed/edited the final manuscript. J.H. performed investigation (co-immunoprecipitation), analyzed the data, visualized the data, and reviewed/edited the final manuscript. J.E.R. performed investigation (CPY* binding and ubiquitination site identification), analyzed the data, visualized the data, and reviewed/edited the final manuscript. J.S. and P.L.F analyzed data (statistical analysis of the DMS dataset) and reviewed/edited the final manuscript. R.D.B. conceptualized the project, performed the investigation, analyzed the data, wrote the original draft, reviewed/edited the final manuscript, and supervised the study.

## Declaration of Interests

The authors declare that they have no competing interests.

## Funding

B.G.P. was supported by the NIH/NIGMS Michigan Predoctoral Training in Genetics (T32GM007544) and NSF Graduate Research Fellowship Program (DGE 1841052). J.E.R. was supported by the NIH/NIGMS Michigan Predoctoral Training in Genetics (T32GM007544). This work was supported by the University of Michigan Medical School Biological Sciences Scholars Program and by NIH/NIGMS Awards (R35GM128592 to R.D.B and R35GM128637 to P.L.F.).

## Supplemental Figure Legends

**Figure S1.**
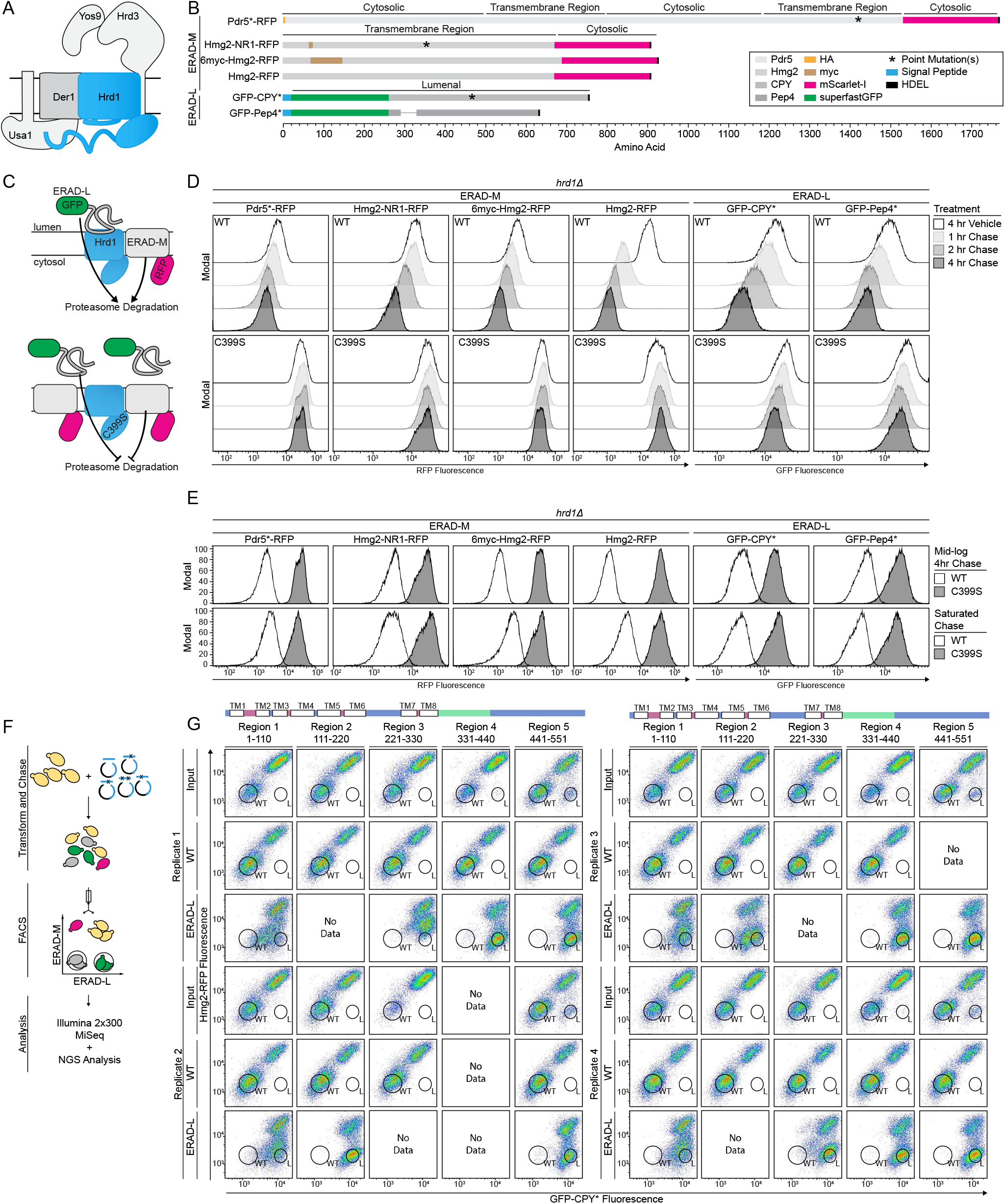
Deep mutation scanning screen development, related to Figure 1. A) Hrd1 complex schematic. B) Schematic of fluorescent ERAD substrates. RFP represents mScarlet-I^58^ and GFP represents superfastGFP^59^. Pdr5* is a mutated ATP-binding cassette transporter with a HA epitope tag inserted between Pro2 and Glu3 that is rendered ERAD-M substrate by a C1427Y mutation^34,60^. There are three different versions of Hmg2 that behave as ERAD-M substrates. Hmg2 represents wild-type Hmg2’s transmembrane region (Met1 to Tyr670) where degradation is regulated by the mevalonate pathway^1^. Hmg2-NR1 contains a myc tag replacing Thr61 through Leu70, and is rendered unresponsive to metabolites in the mevalonate pathway by replacing Thr348 through Ala352 in the sterol sensing domain with a five amino acid substitution (Ile-Leu-Gln-Ala-Ser); this substitution causes Hmg2-NR1 to be constitutively turned over^35^. Finally, 6myc-Hmg2 is a grossly misfolded Hmg2 generated by replacing Ser64 through Glu145 with six tandem myc tags^1^. For ERAD-L substrates, we used CPY* and Pep4*. CPY* is a lumenal vacuolar protease that is rendered a ERAD-L substrate by a single missense mutation of G255R^33^. Pep4* is also a lumenal vacuolar protease that is rendered an ERAD-L substrate by deletion of 37 amino acids from Leu55 through Tyr91^33^. C) Model for fluorescent ERAD reporter proteins. Top: Wild-type Hrd1 degrades fluorescent reporters at the proteasome. Bottom: A RING-finger mutation inactivates Hrd1 (Hrd1(C399S)) causes substrates to accumulate. D) Degradation of the indicated ERAD substrates were followed using flow cytometry after the addition of 0.1% ethanol (vehicle control), 10 µg/mL zaragozic acid (for Hmg2), or 50 µg/mL cycloheximide (for CPY*). Experiments were performed in *hrd1Δ* cells complemented with either wild-type Hrd1(WT) or a RING domain mutant Hrd1(C399S). Histograms are scaled as a percentage of maximum cell count (Modal). E) Substrate degradation was followed using flow cytometry during either mid-log phase growth treated as in (D) (Mid-log Chase, top panels) or with cells grown to saturation and no pharmacological treatment (Saturated Chase, bottom panels). F) Schematic of deep mutational scanning screen. Yeast cells expressing integrated substrates were transformed with a PCR product containing tiling primer mutagenized region of Hrd1 and linearized centromeric plasmid backbone. Transformed cells were grown in liquid culture and subjected to a saturated chase. Wildtype-like cells and ERAD-L defective cells were sorted using FACS. Sorted cells had their phenotype validated before DNA extraction, library preparation, Illumina sequencing, and analysis. During screening optimization, yeast cells expressed substrates from centromeric plasmids and individual colonies were isolated, phenotype validated, and Sanger-sequenced. G) Phenotype confirmation for previously isolated FACS populations following outgrowth (from figure 1D). Top: topology diagram of Hrd1 with transmembrane segments shown as TM1-8. Cytosolic and lumenal segments are shown as blue and magenta respectively. The cytosolic RING domain is shown in green. Bottom: Input library (Input), the wildtype-like sorted population (WT) or ERAD-L defective sorted population were subjected to a saturated chase. The results are displayed as pseudo color flow cytometry plots of GFP-CPY* (x-axis) and Hmg2-RFP (y-axis). Replicate 1 and 2 are on the left set of panels; Replicate 3 and 4 are on the right set of panels. Missing replicates highlighted as “No Data”.

**Figure S2.**
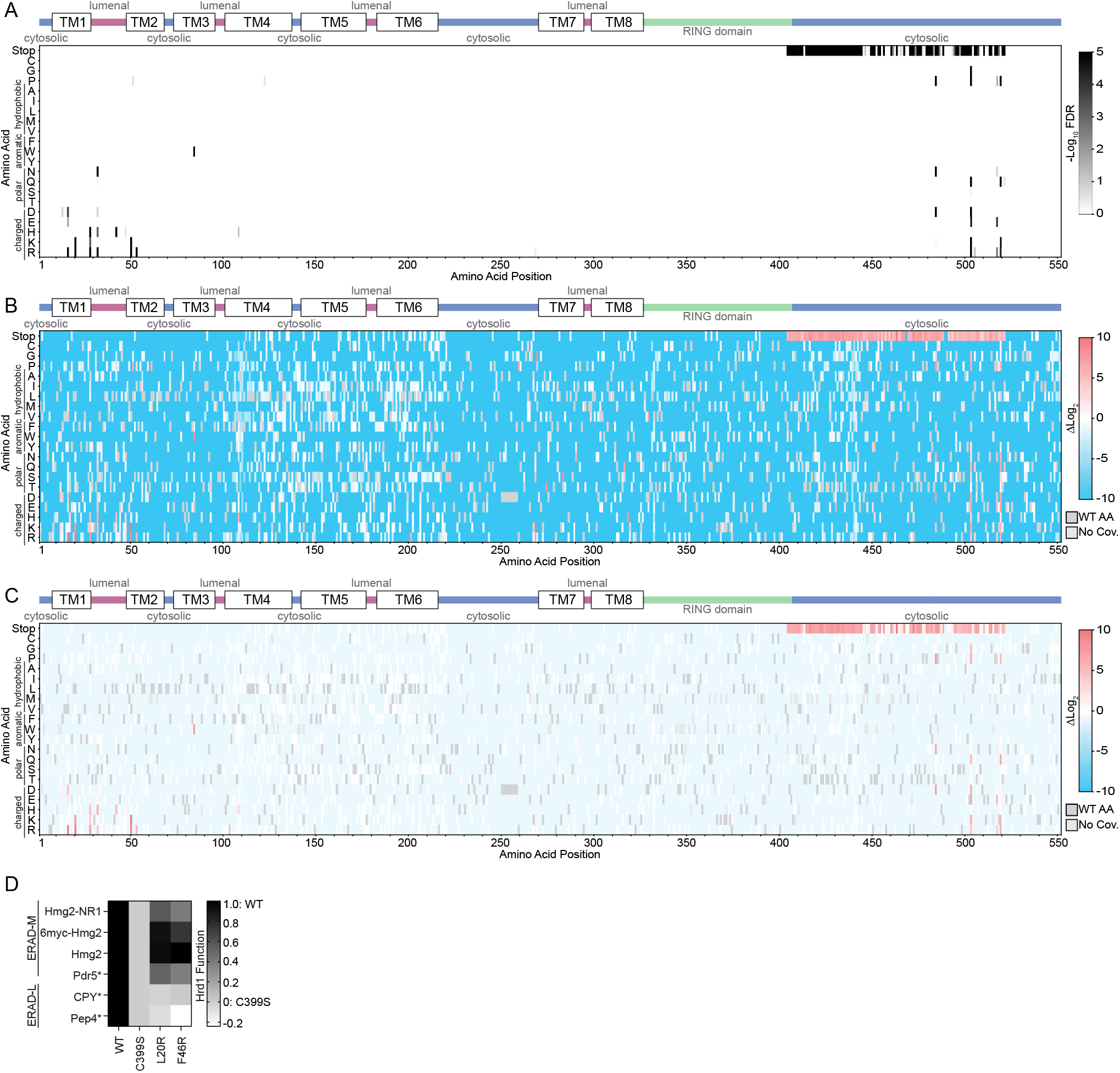
ERAD-L defective DMS results, related to Figure 2. A) Top: Topology diagram of Hrd1 with transmembrane segments shown as TM1-8. Colors indicate the cytosolic (blue), lumenal (magenta), and cytosolic RING domain (green). Bottom: Deep mutational scanning results of cells sorted into the ERAD-L defective bins displayed as a heatmap showing -log_10_(false discovery rate (FDR)). Wildtype amino acids and lack of coverage were omitted for clarity. Related to (Figure 2A). B) As in (A) but showing ERAD-L defective enrichment values. FDR is not used to adjust transparency. C) As in (A) but showing ERAD-L defective enrichment values. Transparency was adjusted based on FDR. FDR below 0.1% were set to 0% transparent and FDR values between 0.1% to 100% were used to adjust transparency from 0% (opaque) to 90% transparent. Individual amino acids are on the y-axis, and the Hrd1 amino acid position is on the x-axis. Dark gray boxes indicate the wildtype amino acid and light gray boxes indicate lack of coverage. D) Degradation of ERAD substrates by individual Hrd1 variants were followed by flow cytometry and summarized in a heatmap. The indicated Hrd1 variants were integrated in a *hrd1Δ* expressing individual ERAD substrates and subjected to a 4-hour mid-log chase. Wild-type Hrd1(WT) is set to 1 (full function) and inactive Hrd1(C399S) is set to 0 (no function). Related to (Figure 2F). For this figure, the number of quantified replicates and individual values are shown in Table S8.

**Figure S3.**
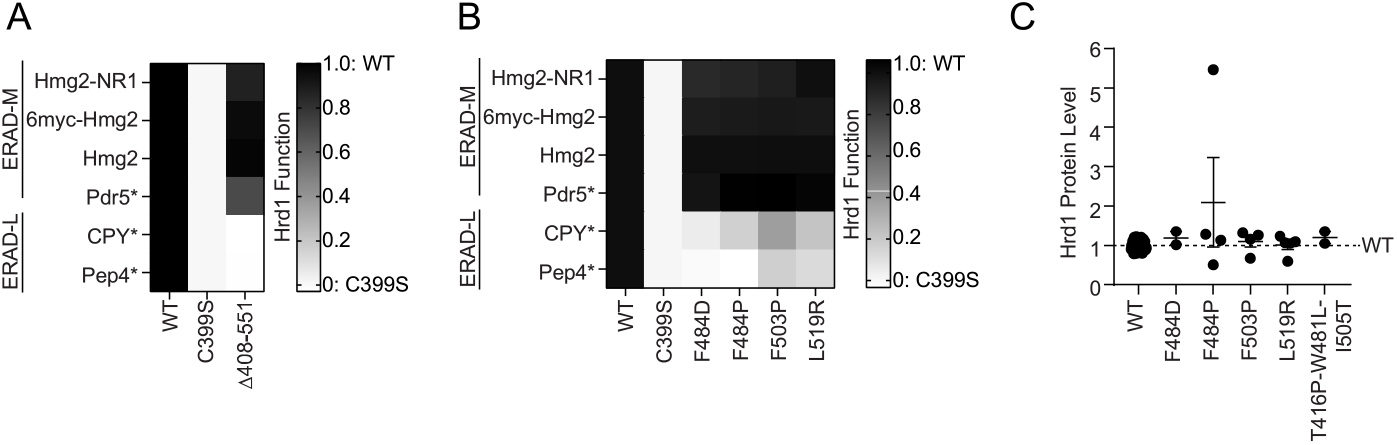
Degradation and protein stability profiles of Hrd1 C-terminal mutants, related to Figure 3. A) Degradation of ERAD substrates by individual Hrd1 variants were followed by flow cytometry and summarized in a heatmap. The indicated Hrd1 variants were integrated in a *hrd1Δ* expressing individual ERAD substrates and subjected to a 4-hour mid-log chase. Wild-type Hrd1(WT) is set to 1 (full function) and inactive Hrd1(C399S) is set to 0 (no function). Related to (Figure 3B). B) As in (A), related to (Figure 3D) C) Expression levels of Hrd1 variants were determined by immunoblotting. *hrd1Δ* cells expressing indicated Hrd1-3xFlag constructs were normalized to wild-type Hrd1(WT) (1-black dash line). Total protein was visualized by stain-free technology as a loading control. Results are displayed as the mean +/− SEM. Individual values are biological replicates. For this figure, the number of quantified replicates and individual values are shown in Table S8 and S9.

**Figure S4.**
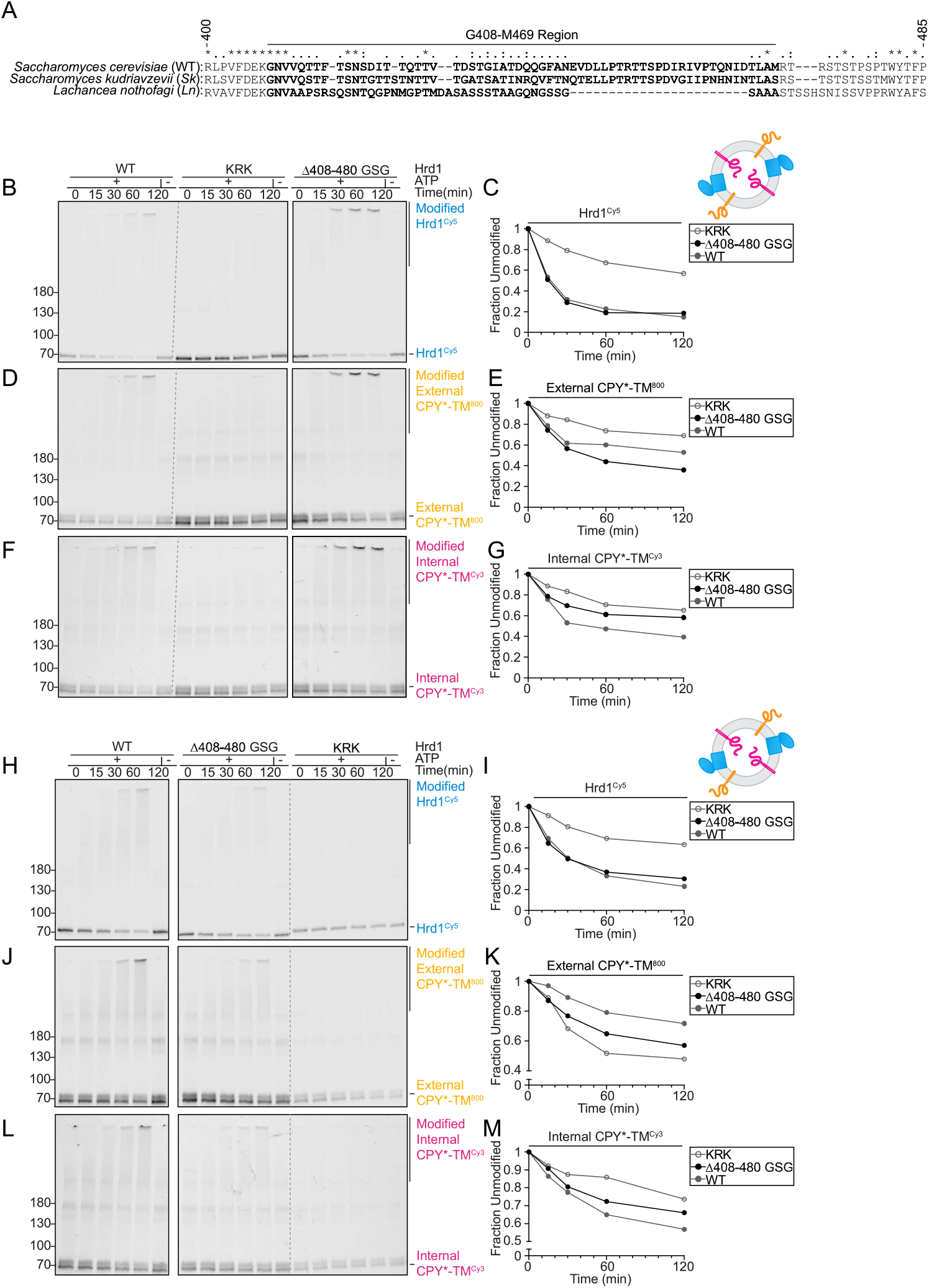
Alignment of divergent Hrd1s and Hrd1(*Δ*408-480_GSG) retrotranslocation, related to Figure 4. A) Sequence alignment (T-Coffee^61^) of Hrd1 from the indicated species. Bold residues highlight the region exchanged to form chimeras used in Figures 4A, 4B, and 4C. B) In vitro autoubiquitination of Hrd1^Cy5^ (blue) in a reconstituted proteoliposome system with externally-oriented CPY*-TM^800^ (orange), internally-oriented CPY*-TM^Cy3^ (magenta). Wild-type Hrd1 (WT-positive control), a retrotranslocation-defective Hrd1 (Hrd1(KRK)-negative control), or Hrd1(*Δ*408-480_GSG) were reconstituted and incubated with recombinant ubiquitination machinery for the indicated times. Samples were analyzed by SDS-PAGE and in-gel fluorescence scanning to visualize Hrd1. Red pixels indicate saturation of signal during the imaging. Related to Figures 4E-4J, second replicate. C) Quantification of (B), showing unmodified Hrd1^Cy5^ (at ∼70 kDa). D) As in (B) showing external CPY*-TM^800^. E) Quantification of (D), showing unmodified external CPY*-TM^800^ (at ∼70 kDa). F) As in (B) showing internal CPY*-TM^Cy^^3^. G) Quantification (F), showing unmodified internally CPY*-TM^Cy3^ (at ∼70 kDa). H) As in (B) showing Hrd1^Cy5^, third replicate. I) Quantification of (H), showing unmodified Hrd1^Cy5^ (at ∼70 kDa). J) As in (H) showing external CPY*-TM^800^. K) Quantification of (J), showing unmodified external CPY*-TM^800^ (at ∼70 kDa). L) As in (H) showing internal CPY*-TM^Cy^^3^. M) Quantification (L), showing unmodified internal CPY*-TM^Cy3^ (at ∼70 kDa).

**Figure S5.**
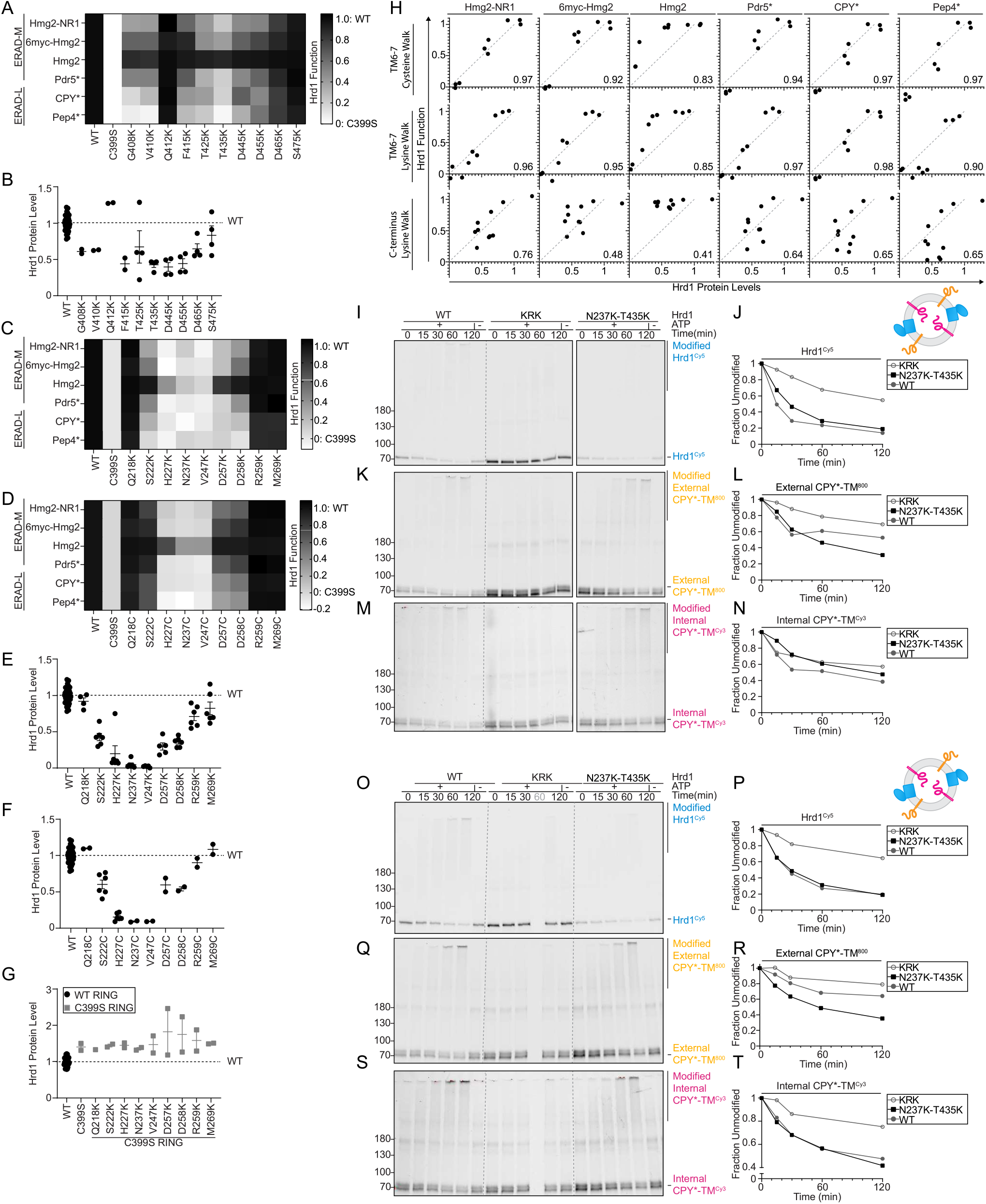
Stability-Function correlation of Hrd1 lysine substitutions, related to Figure 5. A) Degradation of ERAD substrates by individual Hrd1 variants were followed by flow cytometry and summarized in a heatmap. The indicated Hrd1 variants were integrated in a *hrd1Δ* expressing individual ERAD substrates and subjected to a 4-hour mid-log chase. Wild-type Hrd1(WT) is set to 1 (full function) and inactive Hrd1(C399S) is set to 0 (no function). Related to Figure 5C. B) Expression levels of Hrd1 variants were determined by immunoblotting. *hrd1Δ* cells expressing Hrd1-3xFlag constructs from (A) were normalized to wild-type Hrd1(WT) (1-black dash line). Total protein was visualized by stain-free technology as a loading control. Results are displayed as the mean +/− SEM. Individual values are biological replicates. Related to Figure 5D and 5E. C) As in (A) related to (Figure 5G) D) As in (A). E) As in (B) related to (Figure 5H, 5J) F) As in (B) G) As in (B) related to (Figure 5I,5J). Gray values are Hrd1 constructs with the indicated lysine substitutions with the inactivating Hrd1(C399S) RING domain mutation. H) Correlation between Hrd1 protein levels and function for lysine and cysteine substitutions used in (Figures S5A-S5F). Hrd1 protein levels were normalized to wild-type Hrd1 (set to 1) along the x-axis. The y-axis shows Hrd1 function as determined by flow cytometry normalized to inactive Hrd1(C399S) (0) and wild-type Hrd1 (1). Correlation coefficients (Pearson) are displayed in the bottom right corner of each. The gray dashed line demonstrates the trendline for a hypothetical 1:1 ratio of Hrd1 function:protein level. I) In vitro autoubiquitination of Hrd1^Cy5^ (blue) in a reconstituted proteoliposome system with externally-oriented CPY*-TM^800^ (orange), internally-oriented CPY*-TM^Cy3^ (magenta). Wild-type Hrd1 (WT-positive control), a retrotranslocation-defective Hrd1 (Hrd1(KRK)-negative control), or Hrd1(N237K-T435K) were reconstituted and incubated with recombinant ubiquitination machinery for the indicated times. Samples were analyzed by SDS-PAGE and in-gel fluorescence scanning to visualize Hrd1. Red pixels indicate saturation of signal during the imaging. Related to Figures 5M-5R, second replicate. J) Quantification of (I), showing unmodified Hrd1^Cy5^ (at ∼70 kDa). K) As in (I) showing external CPY*-TM^800^. L) Quantification of (K), showing unmodified external CPY*-TM^800^ (at ∼70 kDa). M) As in (I) showing internal CPY*-TM^Cy^^3^. N) Quantification (M), showing unmodified internal CPY*-TM^Cy3^ (at ∼70 kDa). O) As in (I) showing Hrd1^Cy5^, third replicate. Note: Hrd1(KRK) timepoint 60 (gray) was misloaded. P) Quantification of (O), showing unmodified Hrd1^Cy5^ (at ∼70 kDa). Q) As in (O) showing external CPY*-TM^800^. R) Quantification of (Q), showing unmodified external CPY*-TM^800^ (at ∼70 kDa). S) As in (O) showing internal CPY*-TM^Cy^^3^. T) Quantification (S), showing unmodified internal CPY*-TM^Cy3^ (at ∼70 kDa).

**Figure S6.**
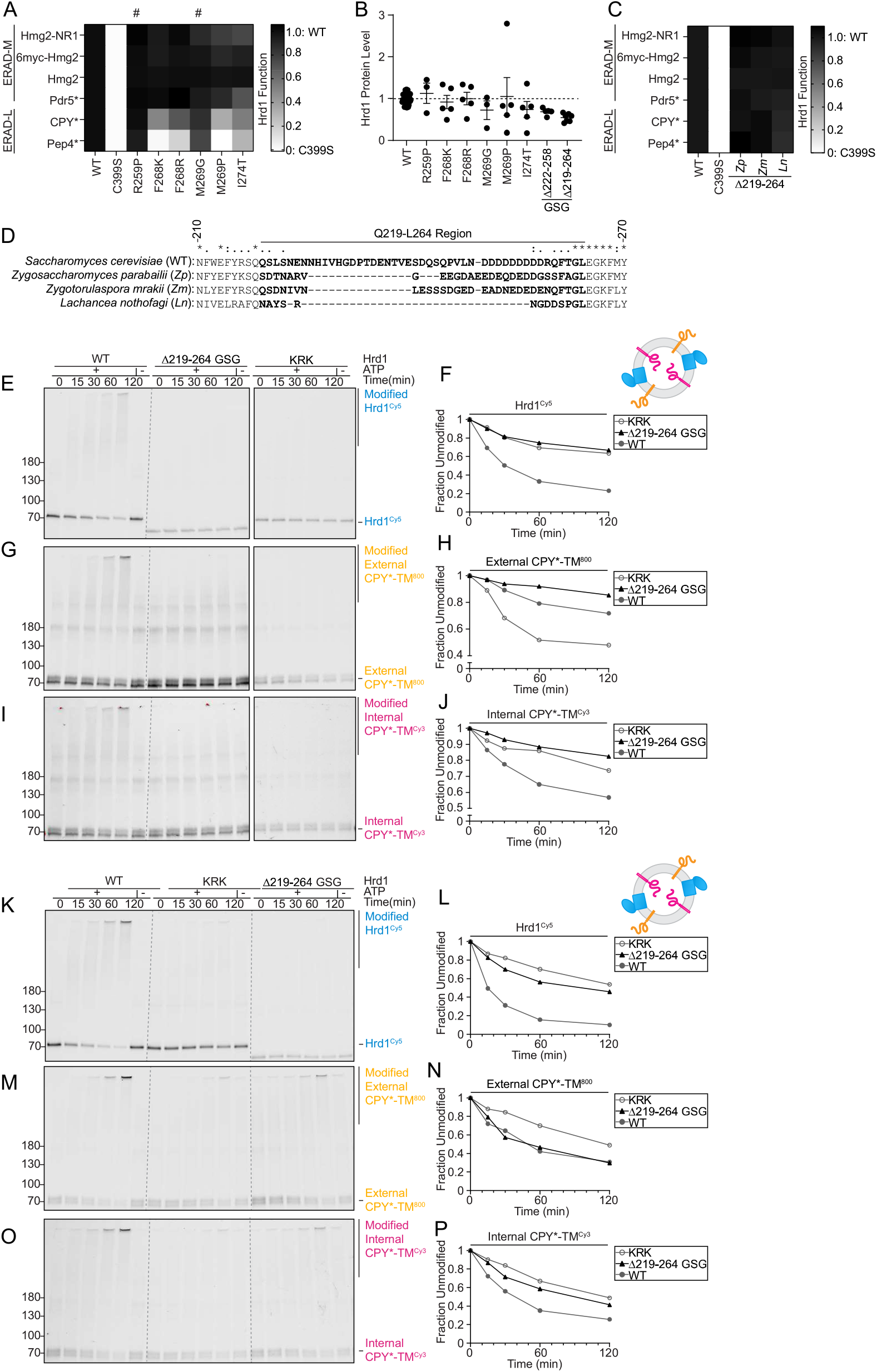
In vivo and in vitro analysis of transmembrane 6-7 loop mutants, related to Figure 6. A) Degradation of ERAD substrates by individual Hrd1 variants were followed by flow cytometry and summarized in a heatmap. The indicated Hrd1 variants were integrated in a *hrd1Δ* expressing individual ERAD substrates and subjected to a 4-hour mid-log chase. Wild-type Hrd1(WT) is set to 1 (full function) and inactive Hrd1 is set to 0 (no function). “#” indicate false positive hits. Related to (Figure 6A). B) Expression levels of Hrd1 variants were determined by immunoblotting. *hrd1Δ* cells expressing indicated Hrd1-3xFlag variants were normalized to wild-type Hrd1(WT) (1-black dash line). Total protein was visualized by stain-free technology as a loading control. Results are displayed as the mean +/− SEM. Individual values are biological replicates. C) As in (A). Related to (Figure 6C). D) Sequence alignment (T-Coffee^61^) of Hrd1 from the indicated species. Bold residues highlight the region exchanged to form chimeras used in Figure 6C and S6C. E) In vitro autoubiquitination of Hrd1^Cy5^ (blue) in a reconstituted proteoliposome system with externally-oriented CPY*-TM^800^ (orange), internally-oriented CPY*-TM^Cy3^ (magenta). Wild-type Hrd1 (WT-positive control), a retrotranslocation-defective Hrd1 (Hrd1(KRK)-negative control), or Hrd1(Δ219-264_3xGSG) were reconstituted and incubated with recombinant ubiquitination machinery for the indicated times. Samples were analyzed by SDS-PAGE and in-gel fluorescence scanning to visualize Hrd1. Red pixels indicate saturation of signal during the imaging. Related to Figures 6E-6J, second replicate. F) Quantification of (E), showing unmodified Hrd1^Cy5^ (at ∼70 kDa). G) As in (E) showing external CPY*-TM^800^. H) Quantification of (G), showing unmodified external CPY*-TM^800^ (at ∼70 kDa). I) As in (E) showing internal CPY*-TM^Cy^^3^. J) Quantification (I), showing unmodified internal CPY*-TM^Cy3^ (at ∼70 kDa). K) As in (E) showing Hrd1^Cy5^, third replicate. L) Quantification of (K), showing unmodified Hrd1^Cy5^ (at ∼70 kDa). M) As in (K) showing external CPY*-TM^800^. N) Quantification of (M), showing unmodified external CPY*-TM^800^ (at ∼70 kDa). O) As in (K) showing internal CPY*-TM^Cy^^3^. P) Quantification (O), showing unmodified internal CPY*-TM^Cy3^ (at ∼70 kDa).

**Figure S7.**
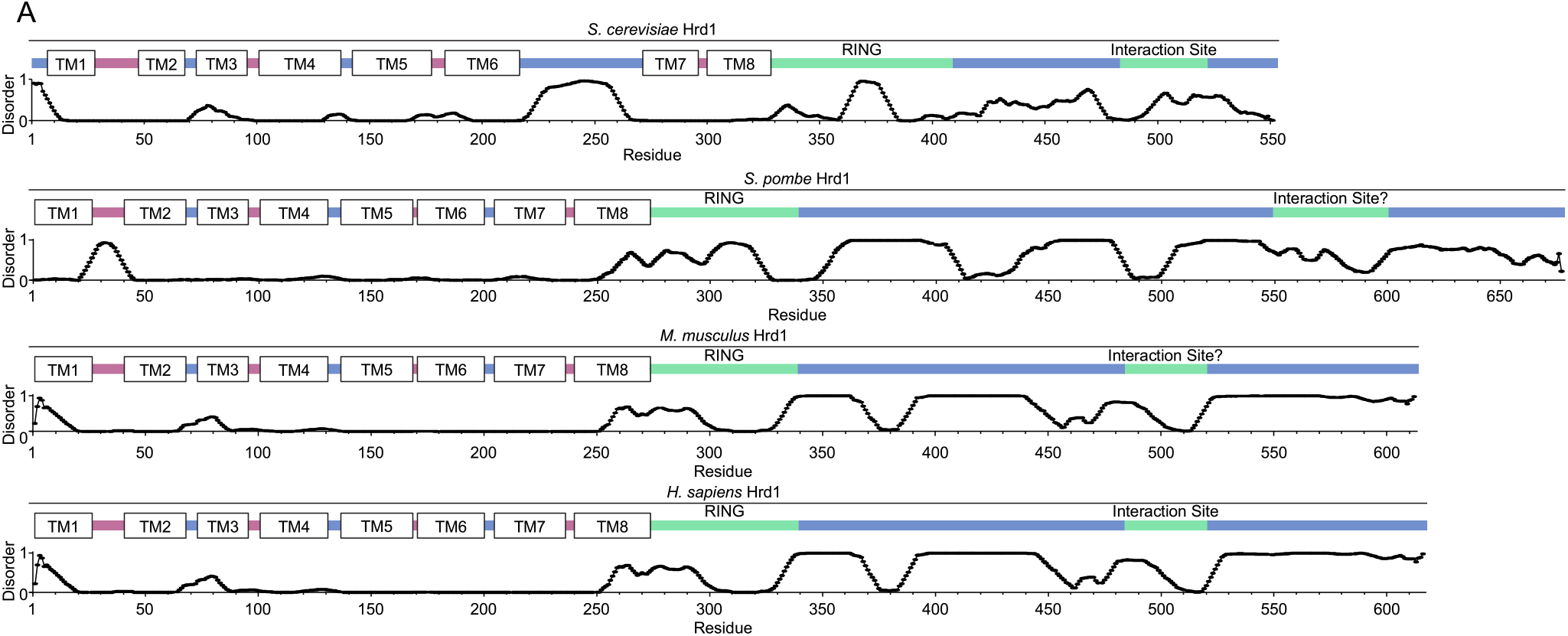
Hrd1 disorder predictions. A) Top: Topology diagram of *S. cerevisiae* Hrd1 with transmembrane segments displayed as TM1-8. Colors indicate the cytosolic (blue), lumenal (magenta), cytosolic RING domain (green), and interaction site (green) is the Usa1 binding site. Bottom: Line chart representing predicted disorder (PONDR VLXT^57^) of *S. cerevisiae* Hrd1 normalized 0 (ordered) to 1 (disordered). B) As in (A) for *S. pombe.* Probable protein interaction site defined by predicted secondary structure in the C-term. C) As in (A) for *M. musculus* Hrd1. Probable protein interaction site inferred from homology of *H. sapiens* Hrd1. D) As in (A) for *H. sapiens* Hrd1. Probable protein interaction site is for HERP and FAM8A1^43^.

## Notes

### Competing Interest Statement

The authors have declared no competing interest.

